# Comparing DNA metabarcoding with morphology in the assessment of macrozoobenthos in Portuguese transitional waters in the scope of the Water Framework Directive monitoring

**DOI:** 10.1101/2022.05.10.491303

**Authors:** Sofia Duarte, Pedro E. Vieira, Barbara R. Leite, Marcos A.L. Teixeira, João M. Neto, Filipe O. Costa

## Abstract

Despite the growing use and potential of DNA metabarcoding to improve and expedite macrozoobenthos monitoring, its employment in Water Framework Directive (WFD) monitoring of transitional ecosystems still remains largely unexplored and pending proof-of-concept studies. In the current study, we addressed this research gap by building upon the biomonitoring network program of the Portuguese Environmental Agency (APA) to benchmark metabarcoding against the morphology-based approach for characterizing macrozoobenthic communities. We assessed the ecological condition of 20 sites from four major transitional ecosystems in the west coast of Portugal, namely Minho, Lima, Vouga and Mondego estuaries. A total of 154 marine invertebrate species were detected with both methodologies, distributed by 11 phyla. In the majority of the sites, metabarcoding returned a higher number of species and phyla than the morphology-based approach (up to 2.5 times higher). In parallel, the proportion of species detected concurrently by both methods was low (35 species, 23%). The use of a multi-locus strategy increased recovered diversity through metabarcoding, since 37 species were detected exclusively with COI and 46 with 18S. For about 61% of the species recovered through morphology, metabarcoding failed detection, among which 20% was due to the lack of reference sequences in genetic databases. For the remaining, we did not find any plausible reason for only 10%, which could be due either to inefficient DNA extraction or PCR failure. Although morphological and metabarcoding-derived biotic indices did not match completely, similar responses to the environmental gradient were obtained in morphology and metabarcoding based-datasets. We anticipate that metabarcoding can increase the throughput and quality of the assessments, allowing faster assessments with greater spatial-temporal density and robust identifications of all specimens in a sample including larval stages, juveniles, and cryptic lineages, as well as smaller taxonomic groups that cannot be identified to species level using the traditional approach.

## 1. Introduction

Coastal and transitional waters are simultaneously among the most important and most threatened ecosystems in the world. They provide important services to humankind, which are at risk due to impactful and diverse anthropogenic pressures (Solan 2004). In the European Union, several legislative measures have been developed aiming to protect and improve the quality of all surface and marine waterbodies, such as the Water Framework Directive (WFD, Directive 2000/60/EC) (European commission 2000) and the Marine Strategy Framework Directive (MSFD, Directive 2008/56/EC) (European commission 2008). WFD requires member states to assess the ecological status of aquatic ecosystems at regular intervals, by sampling or surveying Biological Quality Elements (BQE) following national or EU-wide standard methods. Benthic invertebrates are one of the key BQE employed in WFD, given that they provide invaluable ecosystem services (e.g., support – larval supply, habitat supply; provision – shellfish, genetic resources; regulation –water cleaning and sediments stabilization; cultural – aesthetic), and integrate environmental conditions and changes in a very effective way, which allows the monitoring of long-term responses and site-specific impacts (Salas et al. 2004, Teixeira et al. 2008a,b, Neto et al. 2010, Borja et al. 2011). WFD determines that the macrozoobenthos must be assessed in terms of taxonomic composition, diversity, abundance, disturbance-sensitive and pollution-indicator taxa (European Commission 2011) and several biotic indices have been developed to assess the condition of coastal and transitional waters, with these communities (e.g., Borja et al. 2000, Rosenberg et al. 2004, Teixeira et al. 2009). Some of the most widely used include species richness, Shannon-Wiener, Margalef, AZTI Marine Biotic Index – AMBI, among others, that are assessed through morphology-based identification of specimens (Borja et al. 2000, 2011, Salas et al. 2004, Teixeira et al. 2008b). Resulting measurements are then compared against values expected at “reference conditions”, and water bodies concomitantly assigned to an ecological status (Vinagre et al. 2015).

Although well established and harmonized, bioassessment methodologies of the WFD and MSFD are still intensively debated. Main problems include high monitoring costs and some level of variability and subjectivity that raise unsureness about the reliability of the results. Also, the long time required to complete the species identification process has resulted in the low throughput processing of biomonitoring samples and this is no longer compatible with the need to rapidly reach conclusions about the ecological status of the water bodies (Leese et al. 2018). In addition, despite the imperative need of monitoring and assessment, economic constraints are forcing some countries to reduce the budgets dedicated to biomonitoring (Hering et al. 2018). Due to the above-mentioned reasons, benthic invertebrates monitoring has been conducted most of the times in 1 or 2 events per 6-year management cycle (Hering et al. 2010). This scenario is far from ideal, since benthic invertebrate communities can be susceptible to several natural events that may occur periodically, such as floods and droughts, and that may mask the effects of anthropogenic disturbances and alter ecosystems assessments (Neto et al. 2010). Thus, a more frequent biomonitoring would definitively provide a more accurate and comprehensive view of the present and changing status of benthic ecosystems.

One of the most promising tools for the simultaneous identification of bulk organism assemblages is DNA metabarcoding, where amplicons of standardized DNA-barcode markers, obtained from bulk communities or environmental samples, are massively-parallel sequenced via high-throughput sequencing (HTS) (Hajibabaei 2012, Cristescu 2014). This approach has a number of potential benefits over the morphology-based method, including the simultaneous processing of a large number of samples, increased sensitivity, accuracy and specificity, as well as greater time and cost effectiveness in biodiversity monitoring (Hajibabaei 2012, Cristescu 2014, Duarte et al. 2021).

However, despite the demonstrated utility of metabarcoding to reliably generate assessments of aquatic environmental status, in particular based on benthic invertebrates (Aylagas et al. 2014, 2018, Cowart et al. 2015, Lobo et al. 2017b, Derycke et al. 2021, Duarte et al. 2021, Van den Bulcke et al. 2021), the adoption of metabarcoding in biomonitoring still face several challenges, in particular in coastal and transitional ecosystems. These ecosystems hold very different features from freshwaters, and are highly diverse, thereby requiring a tailored tool in order to overcome existing technological shortcomings that can prevent the detection of all taxa within a sample (e.g., Leite et al. 2021, Wangensteen et al. 2018a,b). In addition, one of the greatest challenges would be to establish a framework for implementation of metabarcoding into monitoring programs, which is currently lacking. To that end it would be imperative to conduct extensive cross-validation studies under realistic scenarios, involving stakeholders, and benchmarking against morphotaxonomic approaches.

In the current study, we evaluated the sensitivity and accuracy of DNA metabarcoding for species detection and identification in the scope of WFD monitoring of macrozoobenthos in coastal and transitional waters in Portugal. To that end we conducted a metabarcoding-morphology comparison on the course of a WFD survey in 20 monitoring sites belonging to four transitional ecosystems – the estuaries of the Rivers Minho, Lima, Vouga and Mondego. This will enable the parallel comparison of morphology and DNA-based data outputs and to test the sensitivity and discriminatory power of metabarcoding on estuarine ecological condition assessment, along selected environmental gradients. To our best knowledge this will be the first attempt in Portugal to address this topic comprehensively (but see Martins et al. 2020, for freshwaters) using metabarcoding, and involving an End-user stakeholder (APA).

## 2. Material and Methods

### 2.1. Sampling sites and environmental characterization

For the current study, 20 sampling sites were selected within 4 estuaries of the west Atlantic coast of Portugal, that are included in the national WFD monitoring program: the estuaries of Rivers Minho (02D/05, 02D/04, 02E/06, 02E/05, 02E/07), Lima (03D/06, 03E/06, 03E/04, 03E/03, 03E/28), Vouga (10E/05, 10E/06, 09F/04, 09F/08, 09E/05, 09F/07) and Mondego (13E/08, 13E/03, 13D/03, 13E/02) and were sampled in late summer 2019 (Fig. 1, please see more details of sampling sites and dates on Supplementary Material: Table S1).

**Figure 1.**
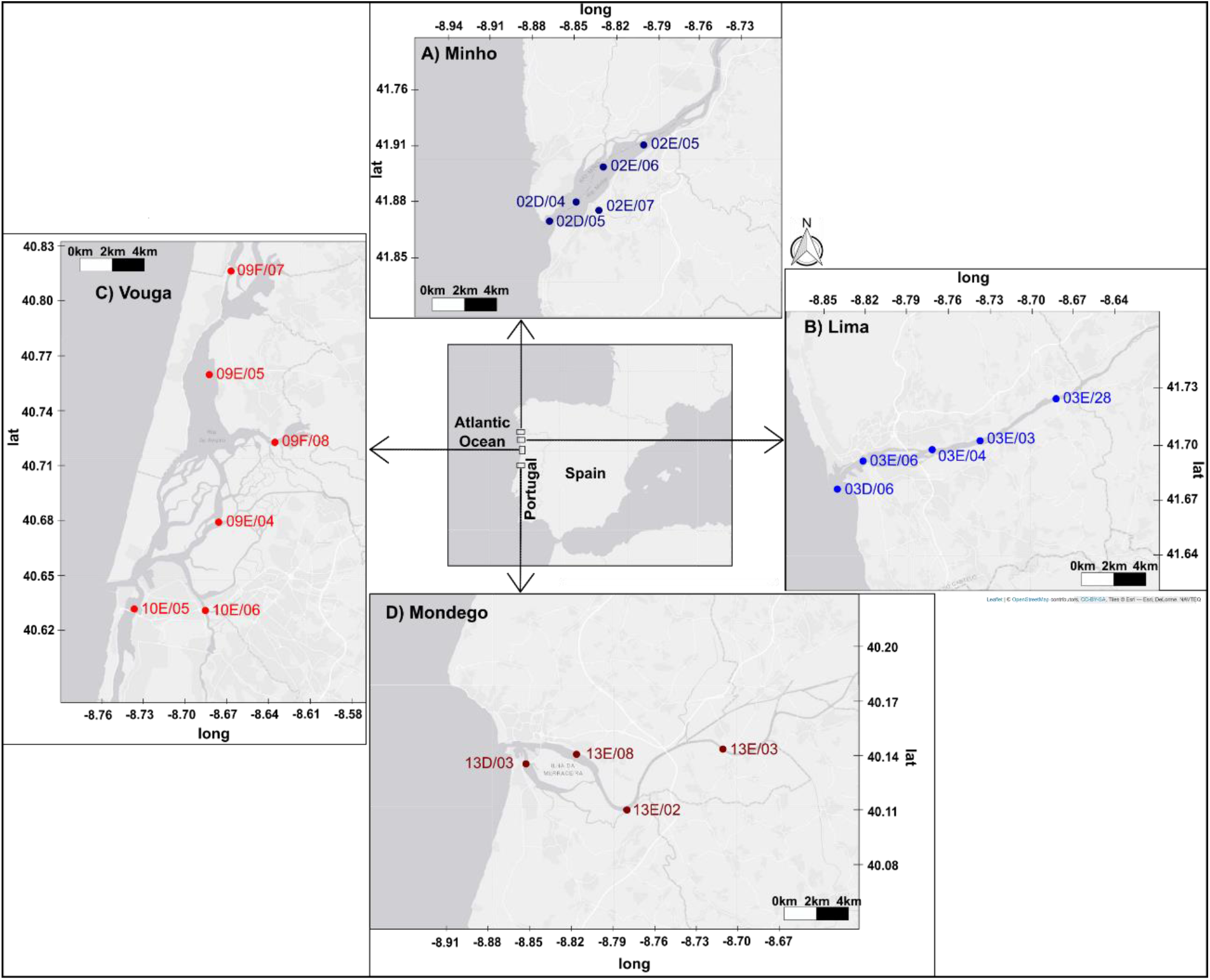
Location of the 20 sites sampled for macrozoobenthos in the scope of the WFD monitoring program, and belonging to four transitional ecosystems: the estuaries of the Rivers Minho, Lima, Vouga and Mondego. See Supplementary Material: Table S1, for more details of the sampling sites.

The Minho estuary is a mesotidal stratified estuary, partially mixed (Sousa et al. 2008) and has been classified as a Natura 2000 site, covering a total estuarine area of 23 km^2^ and located in the Northwest of the Iberian Peninsula. The Lima estuary, is also a mesotidal stratified estuary, partially mixed, and located in the Northwest of the Iberian Peninsula (Sousa et al. 2006, 2007). It is an important harbour in the region, serving trade navigation and fishing activities. Because of this, it has been subjected to constant dredging of the navigation channel within its first 3 km. Other sources of disturbance include the input of agricultural runoff and urban and industrial sewage (Sousa et al. 2007), which changed the physical nature of the lower part of the estuary and have also been responsible for eutrophication (Sousa et al. 2006, 2007). As a consequence of these activities, three of the selected sites are heavily impacted (03D/06, 03E/06 and 03E/04), while the other two remain relatively pristine (sites 03E/03 and 03E/28).

The Vouga estuary is a shallow coastal lagoon and a homogeneous mesotidal estuary, with irregular river discharges, spanning about 75 km^2^ along the central west coast of Iberian Peninsula, where it merges with the freshwater flow of the Vouga River’s catchment area. It has a complex geometry, forming 4 main channels with several branches, exposed to the impact of diverse industrial, shipping, aquaculture and other regional activities (Rodrigues et al. 2011). It is a LTER site (Long Term Ecosystem Research; http://www.lter-europe.net/). Most of the selected sites are pristine (10E/05, 10E/06, 09F/08, 09F/07), but two of them are heavily modified (09E/04, 09E/05). The Mondego estuary is a relatively small warm-temperate homogenous mesotidal system, with irregular river discharges, located in the central west coast of Iberia, formed by two arms; the North and the South arm, which are highly affected by eutrophication (Teixeira et al. 2009). Most of the sites are heavily modified (13E/08, 13E/03, 13E/02), and only one site can be considered pristine (13D/03).

*In situ* water temperature, dissolved oxygen, conductivity, pH and salinity values were registered at high tide conditions, using a YSI professional plus/HANNA HI98194 Ph/EC/DO multiparameter probe. A Niskin/Van Dorn (Horizontal) bottle was used to collect water at the bottom of each sampling site (except for 03E/28, 09F/07 and 13E/03 sites, where water was collected at surface), for total suspended solids (TSS), particulate organic matter (POM), nutrients (NO_3_^-^-N and NH_4_^+^-N) and Silicon (Si) analyses. In the laboratory, the concentrations of NO_3_^-^-N, NH_4_^+^-N and Si were measured using a Skalar San++ Autoanalyser, following adapted and optimised methodologies: NO_3_^-^-N (Houba et al. 1987, Kroon et al. 1993) and NH_4_^+^-N (Krom 1980). Total suspended solids (TSS) were determined by filtering water aliquots, until pre-clogging, through GF/F (0.7 μm pore size, 47 mm diameter, Whatman), previously stuffed and weighed. The filters were then washed with distilled water and dried for 24 hours in a 60 ºC oven, and then weighed to the nearest ± 0.00001 g, after attaining room temperature in a desiccated environment. TSS were assessed, as the difference between dried filters and initial filters weight. For determination of POM, the filters were burned at 450 ºC, for 4 h, and then weighed to the nearest ± 0.00001 g, after attaining room temperature in a desiccated environment. POM was assessed as the difference between dried and burned filters weights. The transparency of water samples was assessed with Secchi disks.

### 2.2. Morphology-based sample processing and taxonomic identifications

Subtidal soft-bottom macrozoobenthic assemblages were sampled with a van Veen grab (sampling area 0.1 m^2^), in the 20 sampling sites of the four transitional ecosystems in late summer of 2019 (for details on sampling dates please see Supplementary Material: Table S1). Three sediment samples were collected from each sampling site, sieved through 1 mm mesh size, and the invertebrate specimens manually sorted from the sediment in the laboratory. Two samples (R1 and R2) were preserved in 4% buffered formalin solution at room temperature and one sample (R3) was preserved in absolute ethanol and placed at 4 ºC until further analysis. Due to logistical reasons only R3 was used for metabarcoding (out of 3 replicates per sampling site). Benthic invertebrates preserved in formalin were counted and identified at the stereomicroscope to the lowest possible taxonomic level, with the assistance of taxonomic identification keys and monographs (e.g., Lincoln 1979, Hayward et al. 1996, Campbell and Nicholls 2008, Hayward and Ryland 2017). Ethanol-preserved bulk invertebrate samples were first subjected to non-destructive DNA extraction, as described below, and subsequently used in morphology-based identification to the lowest possible taxonomic level, as described above.

### 2.3. DNA extraction, preparation of amplicon libraries and high-throughput sequencing (HTS)

Up to 30 g of ethanol-preserved invertebrate samples were used to extract DNA by means of a non-destructive procedure, using a silica-based method, adapted from Ivanova et al. (2006), and as described by Steinke et al. (2022). Briefly, samples were placed in autoclaved flasks, previously washed with 10% bleach and ultra-pure water, to which an adequate volume of a lysis buffer (100 mM NaCl, 50 mM Tris-HCl pH 8.0, 10 mM EDTA pH 8.0 and 0.5% SDS) (depending on the sample wet weight) was added. Samples were then digested overnight in an orbital incubator (Infors) at 140 rpm and 56 ºC. Two aliquots of 1 mL, collected from each lysate, were used in two independent DNA extractions, which were pooled together before PCR amplification. Lysates were then centrifuged and supernatants mixed with a binding mix (6M GuSCN, 20mM EDTA pH 8.0, 10mM Tris-HCl pH 6.4 and 4% Triton X-100) and purified through silica columns and 3 washing steps, with two ethanol-based solutions. DNA was finally eluted from the columns by using autoclaved deionized water. Negative controls were processed along the DNA extraction procedure for checking for contaminations of the solutions used for DNA extractions and labware materials used. These negative controls were used as template in subsequent PCR amplification reactions.

Amplicon libraries and high-throughput sequencing (HTS) were carried out at Genoinseq (Biocant, Portugal). The primer pair mlCOIintF (5’-GGWACWGGWTGAACWGTWTAYCCYCC -3’) (Leray et al. 2013) and LoboR1 (5’-TAAACYTCWGGRTGWCCRAARAAYCA -3’) (Lobo et al. 2013) was used to amplify an internal region of 313 bp of the mitochondrial cytochrome c oxidase I (COI) gene and the primer pair TAReuk454FWD1 (5’-CCAGCASCYGCGGTAATTCC -3’) and TAReukREV3 (5’-ACTTTCGTTCTTGATYRA -3’) (Stoeck et al. 2010, Lejzerowicz et al. 2015) was used to amplify ∼400 bp of the V4 hypervariable region of the 18S rRNA gene (18S). The two primer pairs were selected based on previous studies on marine invertebrates of the region (macrozoobenthos and meiofauna) as the ones that captured the most diverse taxa among four tested primers pairs for COI (Fais et al. 2020, Leite et al. 2021) and three tested primers for 18S (Fais et al. 2020). PCR reactions were performed for each sample using KAPA HiFi HotStart PCR kit according to manufacturer instructions, 0.3 μM of each primer and 50 ng of template DNA, in a total volume of 25 μL. For the mlCOIintF/LoboR1 primer pair, the PCR conditions involved a 3 min denaturation at 95 ºC, followed by 35 cycles of 98 ºC for 20 s, 60 ºC for 30 s and 72 ºC for 30 s and a final extension at 72 ºC for 5 min. For the TAReuk454FWD1/TAReukREV3 primer pair, the PCR conditions involved a 3 min denaturation at 95 ºC, followed by 10 cycles of 98 ºC for 20 s, 57 ºC for 30 s and 72 ºC for 30 s and 25 cycles of 98 ºC for 20 s, 47 ºC for 30 s and 72 ºC for 30s, and a final extension at 72 ºC for 5 min.

Second limited-cycle PCR reactions added indexes and sequencing adapters to both ends of the amplified target regions according to manufacturer recommendations (Illumina 2013). PCR products were then one-step purified and normalized using SequalPrep 96-well plate kit (ThermoFisher Scientific, Waltham, USA) (Comeau et al. 2017), pooled and pair-end sequenced in the Illumina MiSeq® sequencer with the V3 chemistry, according to manufacturer instructions (Illumina, San Diego, CA, USA) at Genoinseq (Biocant, Portugal).

### 2.4. Bioinformatics pipelines

Raw reads, extracted from Illumina MiSeq® System in fastq format, were quality-filtered with PRINSEQ version 0.20.4 (Schmieder and Edwards 2011). This entailed the removal of sequencing adapters and of short reads (<100 bp and <150 bp, for COI and 18S, respectively). Bases with an average quality lower than Q25, in a window of 5 bases were also trimmed. The filtered forward and reverse reads provided by the sequencing facility were merged by overlapping paired-end reads in mothur (make.contigs function, default alignment) (version 1.39.5), where primers sequences were also removed (trim.seqs function, default) (Schloss et al. 2009, Kozich et al. 2013).

The usable reads were then processed in two public databases pipelines (Leite et al. 2021): COI reads were submitted to mBrave – Multiplex Barcode Research and Visualization Environment (www.mbrave.net; Ratnasingham 2019), which is linked with BOLD (Ratnasingham and Hebert 2007) and 18S reads were analysed in SILVAngs database (https://ngs.arb-silva.de/silvangs/; Quast et al. 2013).

In mBrave, COI reads were uploaded using the sample batch function and only the trimming by length was applied (maximum of 313 bp). Low quality reads were then removed if the average quality value (QV) was less than 20 or sequences shorter than 150 bp. Reads fulfilling the previous criteria were further de-replicated and clustered in Operational Taxonomic Units (OTUs) using a distance threshold of 3%. The resultant OTUs were then taxonomically assigned at species level using a 97% similarity threshold against BOLD database that includes several publicly available reference libraries for marine invertebrates of the Northeast Atlantic (e.g., Hollatz et al. 2017, Leite et al. 2020, Vieira et al. 2020).

In SILVAngs, each 18S read was aligned using the SILVA Incremental Aligner (SINA v1.2.10 for ARB SVN (revision 21008)) (Pruesse et al. 2012) against the SILVA SSU rRNA SEED and quality controlled (Quast et al. 2013). Reads shorter than 150 aligned nucleotides and reads with more than 1% ambiguities, or 2% homopolymers, respectively, were excluded from further processing. Putative contaminations and artefacts, reads with a low alignment quality (80 alignment identity, 40 alignment score reported by SINA), were identified and excluded from downstream analysis. After these initial steps of quality control, identical reads were identified (dereplication), the unique reads were clustered (OTUs) on a per sample basis, and the reference read of each OTU was then taxonomically assigned. VSEARCH (version 2.14.2; https://github.com/torognes/vsearch) (Rognes et al. 2016) was used for dereplication and clustering, applying identity criteria of 1.00 and 0.99, respectively. The taxonomic assignment was performed using BLASTn (2.2.30+; http://blast.ncbi.nlm.nih.gov/Blast.cgi) (Camacho et al. 2009) with standard settings and the non-redundant version of the SILVA SSU Ref dataset (release 138.1; http://www.arb-silva.de). The taxonomic classification of each OTU reference read was mapped onto all reads that were assigned to the respective OTU. Reads without any or weak classifications, where the function “(% sequence identity + % alignment coverage)/2” did not exceed the value of 70, remained unclassified and were assigned to “No Taxonomic Match”. In the end, only OTUs taxonomically identified with a similarity threshold of 99% were kept for further analysis.

For both markers, only reads with match at species level were used for further analysis, and taxonomic assignments with less than 9 sequences were discarded (Fais et al. 2020, Leite et al.2021). Any read that matched to non-metazoan and metazoan non-invertebrate groups were also excluded.

In order to maximize the results, the taxonomic assignments were made using the full databases (i.e., BOLD for COI and SILVA for 18S), and each species match was reviewed individually to assess the reliability of the taxonomic assignments. Discordances in the taxonomic assignments were carefully inspected and if they were possible to resolve (i.e., synonyms, clear cases of misidentification), the most probable identification was kept.

### 2.5. Gap-analysis and reasons for the absence of species detection in the metabarcoding dataset

The presence of representative sequences of all the species detected in the present study was assessed in BOLD and SILVA. All the available COI sequences matching the detected species names were mined from BOLD using BAGS (Fontes et al. 2021). All the Animalia records were mined directly from SILVA (version 138.1) to assess which species have representative sequences in this database. A species was considered represented if at least one sequence was available. The reasons for the no-detection of the species in the metabarcoding dataset and that were exclusively identified through morphology were further investigated. Failed detection by one marker or by both may simply have occurred because that particular species was absent in the respective reference databases. However, if a species was present in both reference databases, but was only detected by one marker, or not detected at all through metabarcoding, this would be an indication of inefficient DNA extraction or PCR amplification failure with the primers used for targeting each marker. Other possible reasons pointed out included: 1) species generating less reads than the minimum threshold set in the bioinformatic pipeline (<9 reads); 2) species detection in replicate 1 (R1) and/or 2 (R2), but not in replicate 3 (R3) and 3) inefficient DNA extraction or PCR failure.

### 2.6. Statistical analyses

A principal component analysis (PCA) was used to ordinate sampling sites according to physical and chemical parameters, after data standardization.

Only OTUs with matches to marine invertebrate species (i.e., <3% and <1% ID distance from matching sequences, for COI and 18S, respectively) and with a minimum of 9 reads, were used in further analyses. The number of reads of different OTUs were summed up if they were assigned to the same species, for each marker separately. The validity of the species names for the morphology and the metabarcoding datasets were verified in the World Register of Marine Species (WoRMS) database (WoRMS Editorial Board 2021).

The proportion of species with overlapping or exclusive detections by morphology-based identification and metabarcoding-based identification was determined: 1) for the overall of the species detected, 2) for the species detected within each estuary, and within each estuary 3) for the species detected on each sampling site, using Venn diagrams (http://www.venndiagrams.net/). For each replicate, within each sampling site, the distribution of species among high-rank taxonomic groups (i.e., phyla) was displayed through barplots for the morphology and metabarcoding datasets using GraphPad Prism v6 (GraphPad Software, Inc.).

The AZTI Marine Biotic Index (AMBI) was calculated based on the presence/absence of each species identified through metabarcoding for each marker, and for all species detected using both markers and, based on the presence/absence of each species identified through morphology ((p/a)AMBI) (Aylagas et al. 2014, 2018). This version uses only the presence/absence of each species, ignoring the number of individuals or the number of reads. Values were calculated based on pollution tolerances of the species present, where tolerance is expressed categorically as one of five ecological groups: I, sensitive to pressure; II, indifferent; III, tolerant; IV, second order opportunistic, and V, first order opportunistic (http://ambi.azti.es, Borja et al. 2000) by using the formula: (p/a)AMBI = [(0 x % GI) + (1.5 x % GII) + (3 x %GIII) + (4.5 x % GIV) + (6 x % GV)] / 100; where % represent the number of species falling on each ecological group. (p/a)AMBI index values vary from 0 to 7 indicating different ecosystem status according to pollution tolerances of the species present: 0 to 1.2, unpolluted; 1.3 to 3.3, slightly polluted; 3.4 to 5, moderately polluted; 5.1 to 6, heavily polluted and 6.1 to 7, extremely polluted (Borja et al. 2000, Aylagas et al. 2014, 2018).

Canonical correspondence analysis (CCA) was used to determine the relationships between environmental variables and marine benthic invertebrate’s species detected through morphology and metabarcoding (Ter Braak and Verdonschot 1995). Pearson correlation coefficients were determined to assess the correlations between: i) each environmental variable and PC1 and PC2 axes scores; ii) each environmental variable and CC1 and CC2 axes scores; iii) the no. of specimens (morphology) and the no. of reads (metabarcoding), for species detected with both methodologies in R3; iv) (p/a)AMBI based on morphology assessments and metabarcoding-based assessments; v) (p/a)AMBI and PC1 and PC2 axes scores.

Correlation, PCA and CCA analyses were conducted using PAleontological STatistics (PAST) version 4.03 for Windows (Hammer et al. 2001).

## 3. Results

### 3.1. Environmental characterization of the sampling sites within each estuarine system

The full environmental characterization of each sampling site within each estuary is detailed in Supplementary Material: Table S1 and summarized in the PCA diagram displayed in Fig. 2.

**Figure 2.**
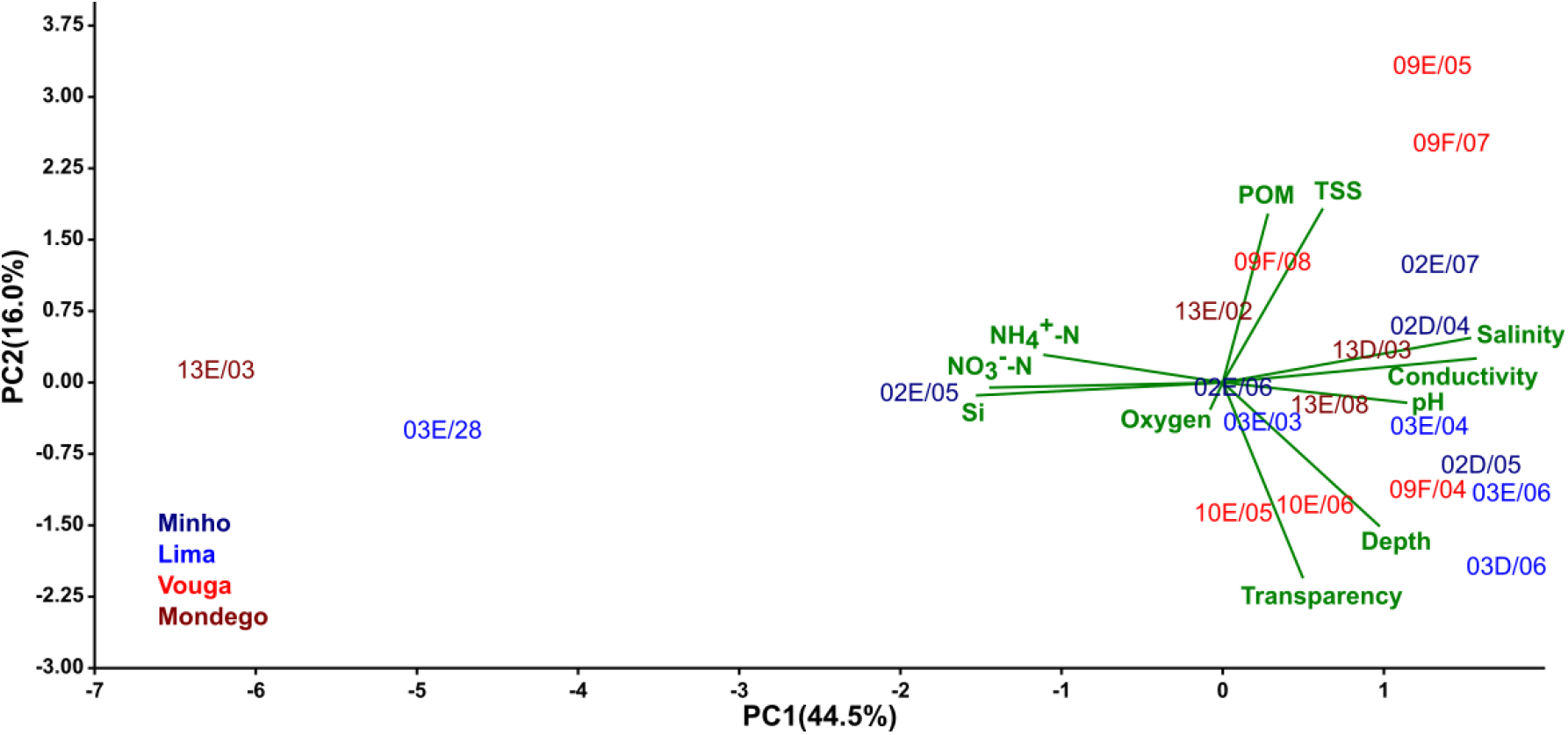
Principal components analysis (PCA) of physical and chemical stream water parameters of the 20 sampling sites along the 4 estuaries.

PCA ordination of the 20 sites according to water variables showed that axes 1 and 2 explained 60.5% of the total variance (Fig. 2). The first axis was significantly correlated with salinity, conductivity and pH, and nitrates, ammonia and Si concentrations which changed in opposite directions (more details of the coefficients of correlation and p values on Supplementary Material: Table S2). The second axis was more correlated with TSS and POM, and transparency and depth, which also changed in opposite directions (Supplementary Material: Table S2). There was a clear separation of the oligohaline sites 13E/03 (Mondego) and 03E/28 (Lima) from the remaining sites, which according to the available salinity values were classified as poly-mesohaline (Minho - 02E/06, 02E/05; Lima - 03E/03; Mondego – 13E/02) and euhaline (Minho - 02D/05, 02D/04, 02E/07; Lima - 03D/06, 03E/06, 03E/04; Vouga - 09F/08, 09E/05, 09F/07; Mondego - 13E/08).

### 3.2. Initial metabarcoding dataset processing

The number of initial raw reads was 1111837 and 770841 for COI and 18S, respectively (Table 1). Subsequent filtering steps (short length reads removal, de-multiplexing, primers removal, de-replication and chimera’s removal) reduced the number of sequences to 718120 and 448299 for COI and 18S, respectively (Table 1). From these, 663364 and 448291 were taxonomically classified for COI and 18S (Supplementary Material: Table S3), respectively, and from these 55.4% and 38.8%, of the initial reads, matched with marine invertebrate taxa, while 52.5% and 37.3% had species level assignments with sequence numbers superior to 8, for COI and 18S, respectively (Table 1). The % of COI reads that matched marine invertebrate species varied between 2.4% and 85.1%, for 13D/03 (Mondego) and 02E/06 (Minho), respectively, while the % of 18S reads that matched marine invertebrate species varied between 0 and 64.2%, for 02E/06 (Minho) and 03D/06 (Lima), respectively (Supplementary Material: Table S3).

**Table 1.** Total number of sequences generated in Illumina MiSeq high-throughput sequencing (raw reads) and %, in comparison with initial raw reads, retained after all the processing steps of the bioinformatics pipeline (demultiplexing, primers removal and quality filter) (usable reads), and assigned to marine invertebrate taxa and species, for each marker (COI and 18S). *, reads submitted to taxonomic assignment; **, reads with taxonomic classification at species level (>97% for COI and >99% for 18S) and sequence number higher than 8.

|  | Marker |  |
| --- | --- | --- |
|  | COI | 18S |
| Raw reads | 1111837 (100%) | 770841 (100%) |
| Usable sequences* | 718120 (64.6%) | 448299 (58.2%) |
| Taxonomic match w/ marine invertebrates | 615521 (55.4%) | 299082 (38.8%) |
| Species level taxonomic assignment (>8 reads)** | 584220 (52.5%) | 287397 (37.3%) |

### 3.3. Morphology and metabarcoding-based benthic invertebrates’ taxonomic assignments

A total number of 154 marine invertebrate taxa was identified at species level, by using both methodologies (Fig. 3, Supplementary Material: Table S4). From these, 90 species were identified through morphology, while 99 species through metabarcoding (Fig. 3, Supplementary Material: Table S4). Thirty-five species were identified by using both methodologies (ca. 23%), while 55 species were exclusively detected through morphology (ca. 36%) and 64 through metabarcoding (ca. 42%) (Fig. 3A, Supplementary Material: Table S4).

**Figure 3.**
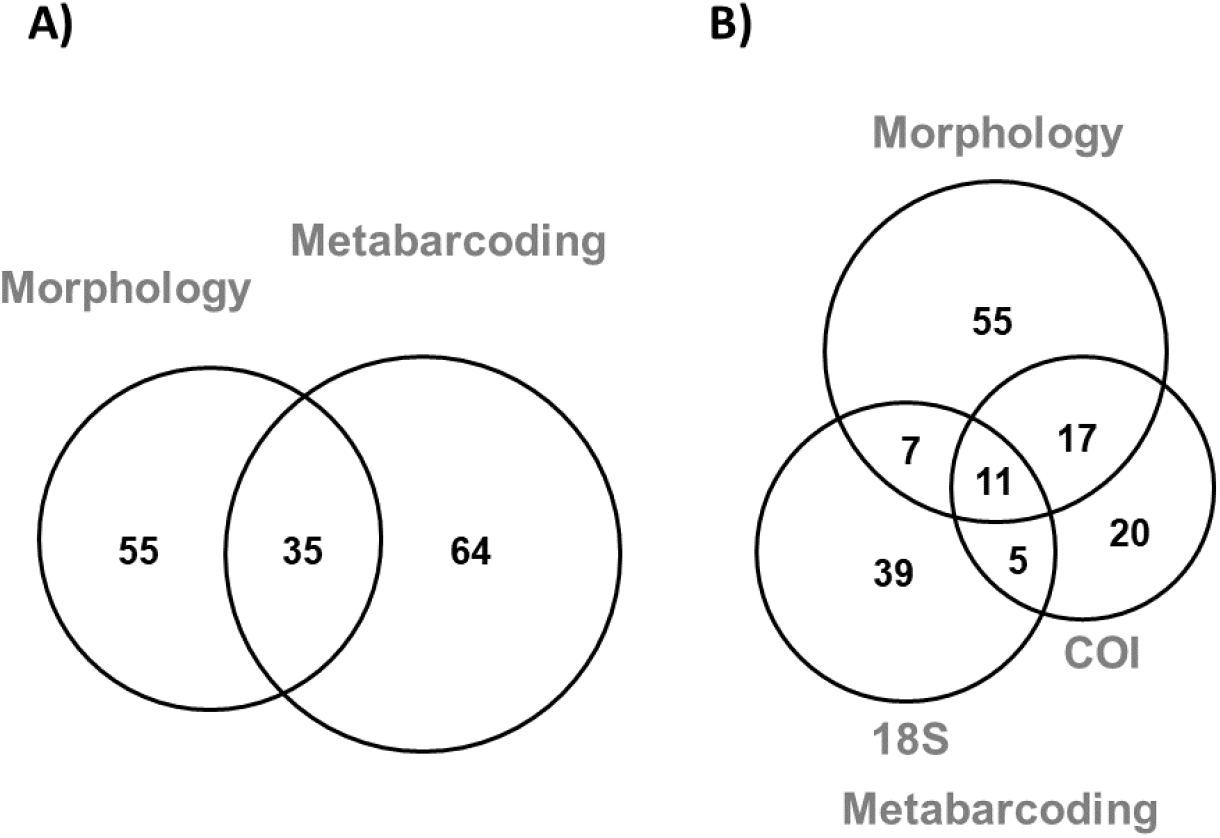
Partitioning of the marine invertebrate species detected with both methodologies (A) and detected with both methodologies, but discriminated by genetic marker for the metabarcoding dataset (B). All sampling sites were considered.

Within the metabarcoding dataset, 16 species were detected with both markers (ca. 16%), while 37 species were exclusively detected with COI (ca. 37%) and 46 with 18S (ca. 47%) (Fig. 3B, Supplementary Material: Table S4).

The highest numbers of total marine invertebrate species were found in Lima (80 species) and Vouga estuaries (81 species), while lower numbers were recovered from Minho (35 species) and Mondego estuaries (41 species) (Fig. 4). In general, metabarcoding recovered equal or higher diversity of species than morphology-based identification in the more diverse estuaries (Lima and Vouga), while the opposite was found for the lowest diverse estuaries (Minho and Mondego). The % of species recovered with both methodologies varied between 17% and 25%, for Vouga and Lima estuaries, respectively. A closer look into each sampling site revealed that for about half of the sites, metabarcoding recovered a higher diversity of species than morphology-based identification (Supplementary Material: Fig. S1) and the % of the diversity recovered with both methodologies varied between 0 and 20%, for Minho; 6 to 40%, for Lima; 5 to 36%, for Vouga and 0 to 27%, for Mondego, respectively (Supplementary Material: Fig. S1).

**Figure 4.**
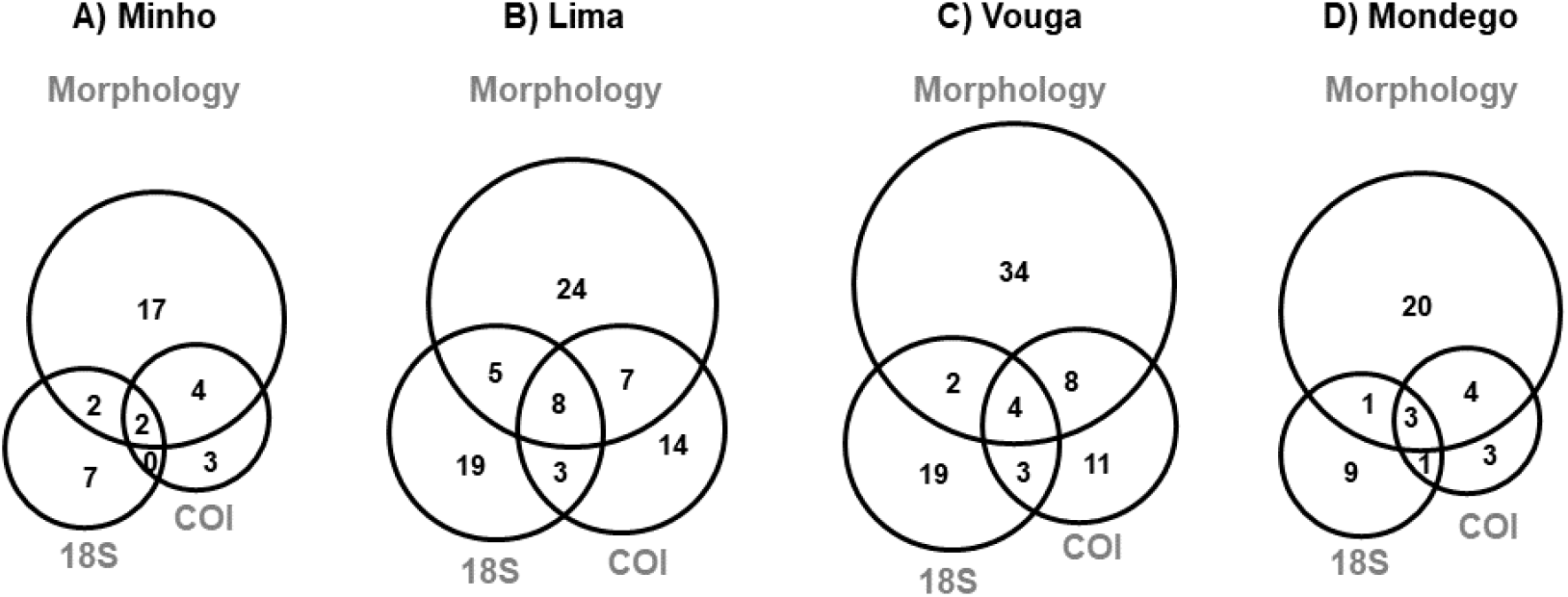
Partitioning of the marine invertebrate species detected with both methodologies for Minho (A), Lima (B), Vouga (C) and Mondego (D) estuaries. All sampling sites were considered, within each estuary.

A closer look within each estuary indicated that maximum numbers of marine invertebrate species were always detected with metabarcoding for all sampling sites in Lima and almost all in Vouga estuaries (except 09E/05) (maximum of 30 in 03D/06 and 21 in 09E/05, for Lima and Vouga, respectively) (Fig. 5). In Minho estuary, while higher numbers of species were retrieved through metabarcoding in 3 out of 5 sampling sites (02D/05, 02D/04, 02E/05), a maximum number of 9 species was found through morphology-based identification in 02E/07. In Mondego, a higher number of species was also retrieved through metabarcoding in 2 out 4 sampling sites (13E/08 and 13E/03), but the highest diversity retrieved (15 species) was attained in 13D/03, through morphology-based identification.

**Figure 5.**
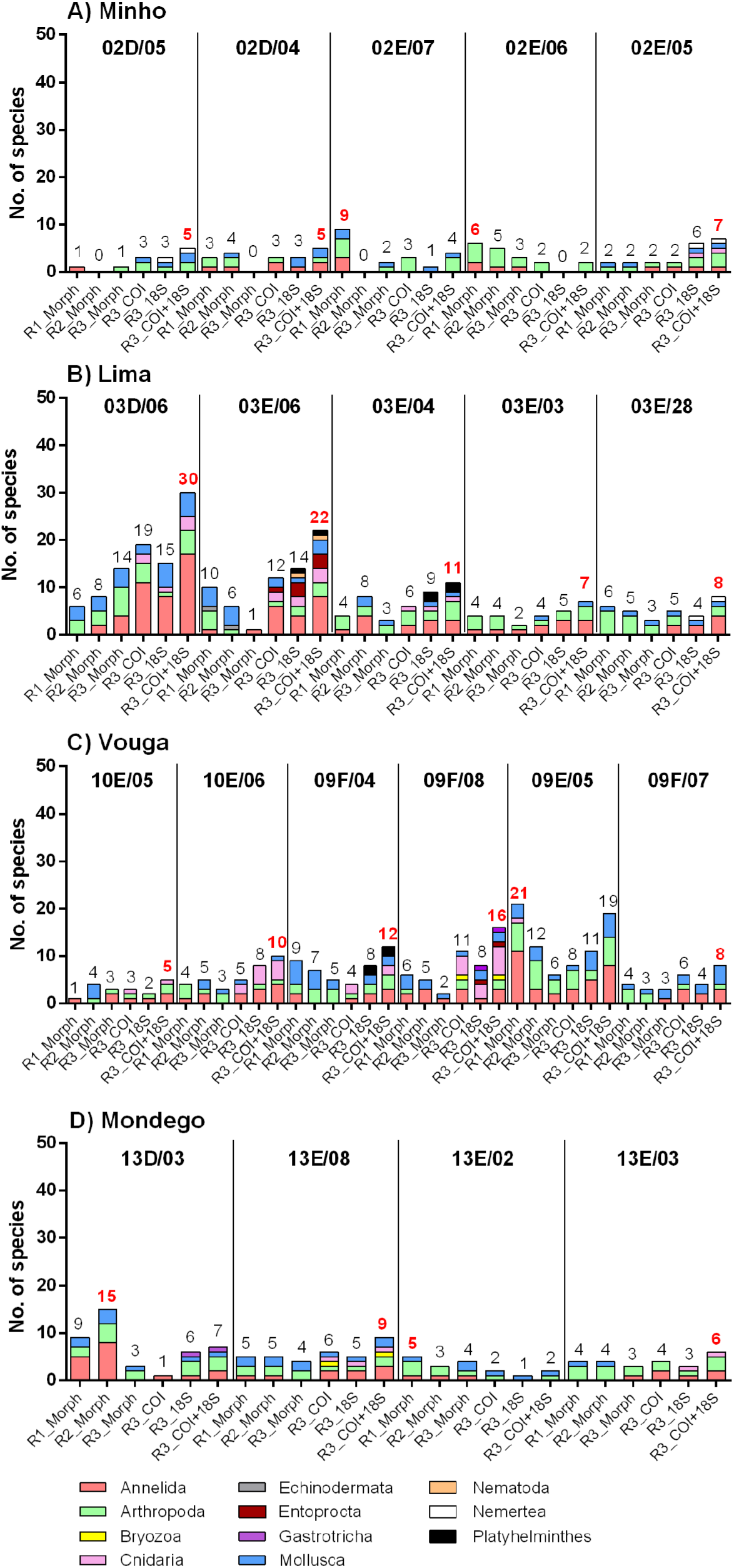
Taxonomic profiles of marine invertebrate species detected through morphology and metabarcoding (COI, 18S and COI + 18S) on each analysed replicate collected on each sampling site on Minho (A), Lima (B), Vouga (C) and Mondego (D) estuaries. Numbers above bars indicate the total number of species detected on each sampled replicate (in red is represented the highest number of species detected on each sampling site).

The 154 marine invertebrate taxa identified at species level were distributed by 11 taxonomic groups: Annelida, Arthropoda, Bryozoa, Cnidaria, Echinodermata, Entoprocta, Gastrotricha, Mollusca, Nematoda, Nemertea and Platyhelminthes (more details of the species names and the associated taxonomic groups in Supplementary Material: Table S4). Echinodermata species were exclusively detected through morphology, but not by metabarcoding, while Bryozoa, Entoprocta, Gastrotricha, Nematoda, Nemertea and Platyhelminthes were exclusively detected through metabarcoding, with the 4 later being exclusively detected with the 18S marker (Fig. 5). In general, the most well represented groups included Arthropoda (42.2%), Annelida (33.3%) and Mollusca (22.2%), in the morphology dataset, and Annelida (42.3% and 27.4%), Arthropoda (28.8% and 14.5%), Mollusca (13.5% and 17.7%) and Cnidaria (13.5% and 16.1%, for COI and 18S, respectively), in the metabarcoding datasets. Cnidaria, were particularly abundant in Vouga estuary, more specifically in sampling sites 10E/06 and 09F/08.

Since 55 invertebrate species (out of a total of 90, 61%) were exclusively detected through morphology (Supplementary Material: Table S5), we further investigated the reasons for this. From this list, 18 species do not have sequences belonging to any of the targeted markers on genetic databases (20%) (Fig. 6A, Supplementary Material: Table S5). A closer look revealed that 26 species were detected in replicates R1 and/or R2, but not in R3, the replicate analysed through metabarcoding (Fig. 6B). Other reason included 2 species producing less reads than the minimum threshold set in the bioinformatic pipeline (<9 reads). For the remaining 9 species (10%) the absence of detection in the metabarcoding dataset could be due either to inefficient DNA extraction or PCR amplification failure with the primers used for targeting each marker.

**Figure 6.**
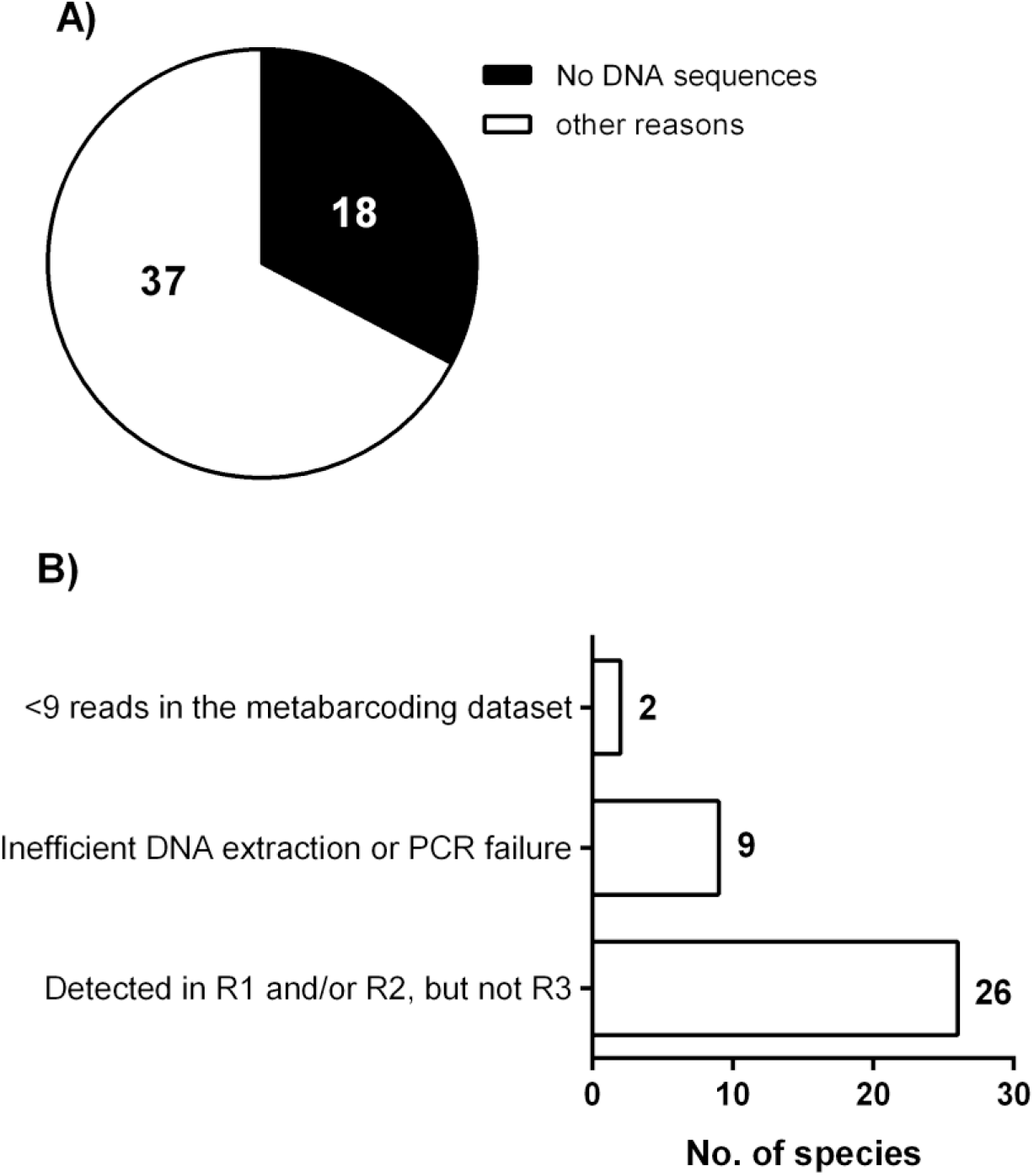
Number of marine invertebrate species exclusively detected with the morphology-based approach, represented and not represented with DNA sequences in genetic databases (COI and/or 18S) (A) and possible reasons for the no detection through metabarcoding for species that were represented with DNA sequences in genetic databases (B).

### 3.4. Morphology and metabarcoding-derived biotic indices

The % of each ecological group, based on marine invertebrate species, varied across sampling sites, and slight differences were found between both methodologies (Fig. 7). Since no clear relationship was found between the number of reads and the number of specimens found, for each species detected with both methodologies (Supplementary Material: Table S6, Fig. S2), we opted to calculate the % of each ecological group and AMBI indices based solely on the absence/presence of species ((p/a)AMBI).

**Figure 7.**
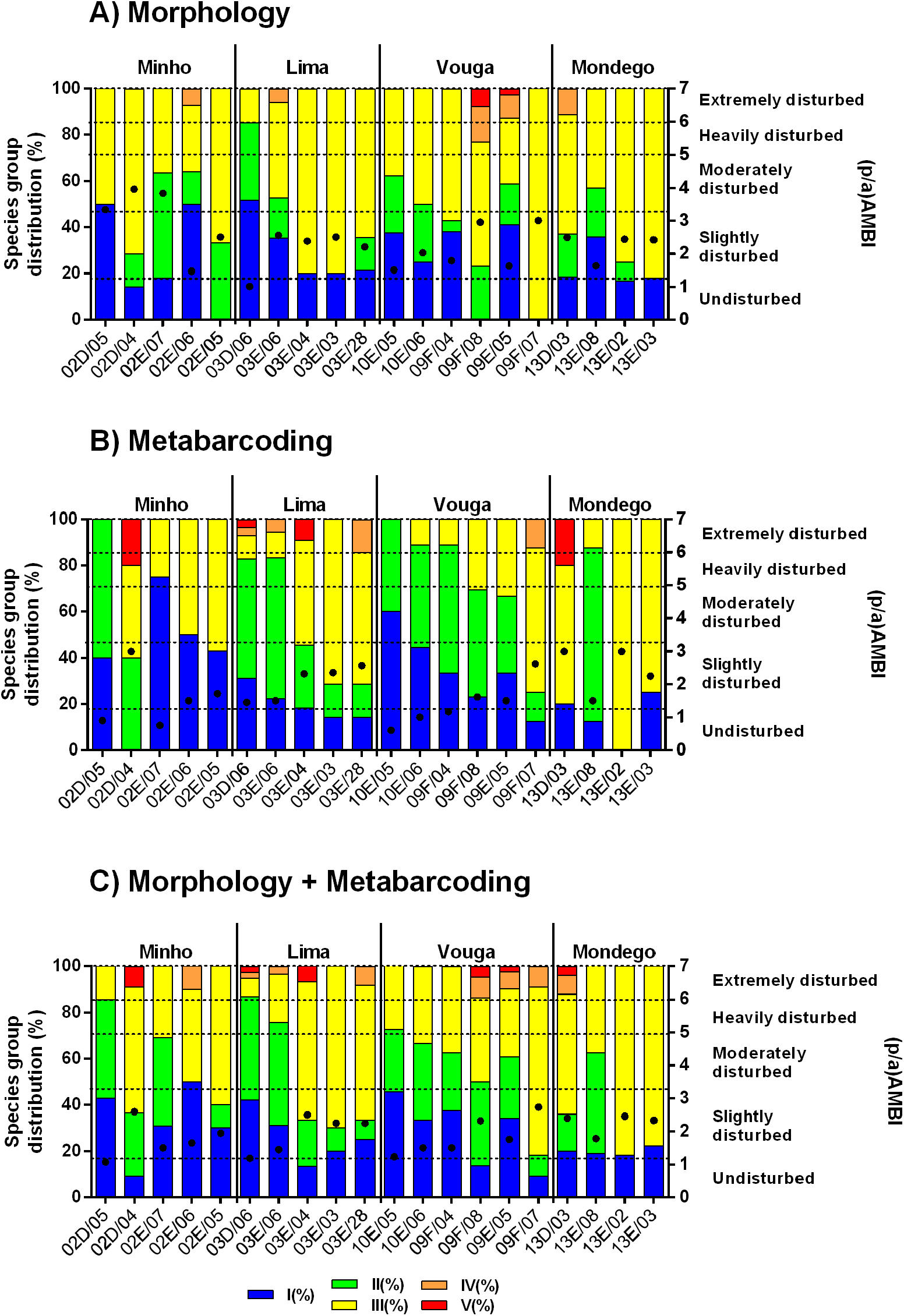
% contribution of each ecological group at each sampling site based on presence and absence of each marine invertebrate species identified using morphology (A), metabarcoding (B) and the combination of both approaches (C). Dots on each bar represent (p/a)AMBI indices scores obtained at each sampling site.

By using morphology-based identification (Fig. 7A), class III ecological group dominated in most sampling sites (in 15 out of 20), resulting in most of the sites being attributed with a classification of slightly disturbed, with the exception of 03D/06 from Lima, which was classified as undisturbed, and 02D/05, 02D/04 and 02E/07 from Minho, which were classified as moderately disturbed.

For the metabarcoding dataset (Fig. 7B), the distribution of the species among the different ecological groups was more variable. Even so, class III ecological group dominated or co-dominated in 11 of the sampled sites, while in the other 9, the dominant or co-dominant ecological groups included class II (10 out of 20) and to a lesser extent class I (4 out of 20) ecological groups. Most of the sampling sites were still classified as slightly disturbed (15), while 5 sites as undisturbed (02D/05 and 02E/07 in Minho and 10E/05, 10E/06 and 09F/04 in Vouga). (p/a)AMBI values calculated using morphology and metabarcoding-based assessments were poorly correlated (Supplementary Material: Fig. S3), probably due to the differences in the % of the ecological groups detected between both methodologies. When joining the two datasets (Fig. 7C), class III ecological group still dominated or co-dominated in most of the sampling sites (11), class II in 7 sites and class I in 6 sites. All sampling sites were classified as slightly disturbed, with the exception of sites 02D/05, in Minho, and 03D/06, in Lima, which were classified as undisturbed. No significant correlations were found between (p/a)AMBI indices and PC1 and PC2 axes scores of environmental variables from Fig. 2, with the exception of (p/a)AMBI values of the combined datasets which were significantly correlated with PC2 (Supplementary Material: Fig. S4).

Canonical correspondence analysis of the relationships between physical and chemical parameters of the water and the structure of marine invertebrate species based on morphology (Fig. 8A), metabarcoding (Fig. 8B) and in the combination of both approaches (Fig. 8C), showed that the first two axes explained ca. 32%, 30% and 30% of the total variance, respectively. For both datasets (morphology and metabarcoding) and when combined, there was a clear separation of invertebrate communities from the oligohaline site 03E/28 (Lima), from the remaining sites. The distribution of the sites accordingly to the invertebrate community structure derived from morphology, metabarcoding and from the combination of both approaches was very similar along the environmental gradient defined by axis 1 and axis 2. Despite the small differences found, in all datasets, the variables that most significantly affected community structure were salinity, conductivity and Si (details of r coefficients and p values in Supplementary Material: Table S7). Other variables that significantly affected community structure included transparency, maximum depth and nitrates concentrations (Supplementary Material: Table S7).

**Figure 8.**
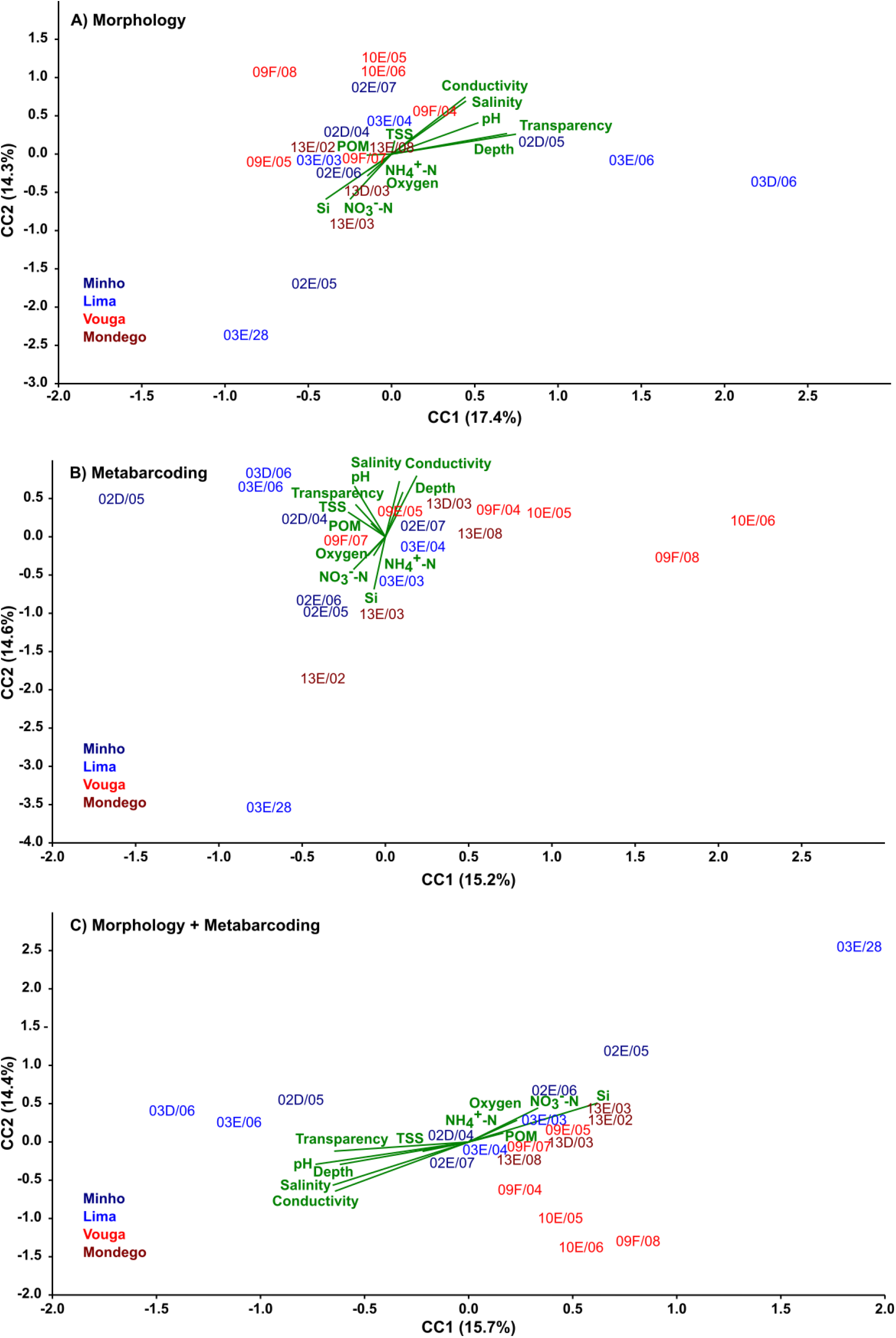
Canonical correspondence analysis diagrams for the ordination of physical and chemical parameters and invertebrate species at the 20 sampled sites, based on morphology (A), metabarcoding (B) and morphology plus metabarcoding (C).

## 4. Discussion

Although the use of DNA metabarcoding to monitor aquatic benthic invertebrate communities has been growing steeply in the last few years (Duarte et al. 2021), few studies targeted transitional ecosystems (Chariton et al. 2015, Lobo et al. 2017b, Aylagas et al. 2018, Steyaert et al. 2020); therefore, they require further and more exhaustive proof-of-concept studies. In the current study, we addressed this research gap by building upon the WFD biomonitoring surveys of the Portuguese Environmental Agency (APA), to assess the ecological condition of 20 sites from four main transitional ecosystems in the west coast of Portugal – the estuaries of Lima, Minho, Vouga and Mondego Rivers. In fact, at least to our knowledge, only one previous study involved an official biomonitoring program to benchmark metabarcoding against the morphology-based approach for characterizing benthic invertebrate communities in transitional ecosystems (Aylagas et al. 2018, in the Basque country, Spain). In Portugal, this is the first time that a study of this kind is conducted in transitional waters, but see Martins et al. (2020) for freshwater ecosystems.

### 4.1. Metabarcoding outperformed morphology-based assessments of benthic invertebrate species

Compared to the morphology-based approach, metabarcoding was able to recover an equal or higher species richness and phylogenetic diversity in 10 out of the 20 sampling sites (Supplementary Material: Fig. S1). The significance of this result is reinforced by the fact that this greater species detection ability was accomplished using only one replicate in metabarcoding, compared to 3 replicates in morphology-based assessments. However, when comparing exactly the same replicate, where both approaches were concurrently employed, metabarcoding always outperformed the morphology-based approach, with the exception of 2 sampling sites (02E/06 and 13E/02), where the differences were minute (3 and 4 *versus* 2 species, for morphology and metabarcoding, respectively) (Fig. 5). In some cases, the differences were fairly high, the most notable for sampling site 03E/06, where a total of 22 species were recovered through metabarcoding (12 and 14, for COI and 18S, respectively), in comparison to only one species detected through morphology (the polychaete trumped worm *Lagis koreni*). Several metabarcoding studies have reported the occasional detection of species that were apparently lacking through the morphological analyses of the same samples (Cowart et al. 2015, Aylagas et al. 2016, 2018, Hollatz et al. 2017, Lobo et al. 2017b, Cahill et al. 2018, Steyaert et al. 2020). Although organisms were carefully sorted from the debris before DNA extraction, eventually small portions of associated inorganic (sediment) and organic detritus (algae) were also picked. These may contain tiny organisms, body fragments or tissues, and even gut contents or DNA from ecologically associated species that can be detected through metabarcoding, due to the high sensitivity of the technique (Aylagas et al. 2016, Hollatz et al. 2017, Lobo et al. 2017b). In fact, a closer look into the diversity found in this particular sample (03E/06) revealed the presence of three hydrozoans, three entoprocts, one nematode and one platyhelminth species in the metabarcoding dataset, which could have been easily unnoticed through the morphological inspection. Interestingly, although undetected through morphology, the DNA of *Abra alba* was also detected in this sample, a species that is often reported to live in close association with *Lagis koreni* (Thiébaut et al. 1997, Bacouillard et al. 2020). Similarly, *Lagis koreni* was detected only through metabarcoding in a sample (03D/06) where high densities of *Abra alba* were recorded by morphological inspection.

In addition, the fraction of species detected by both methods was quite low: 23% if we considered all the dataset (Fig. 3); 17% to 25%, at the estuary level and 0 to 40%, at site level (Fig. 4 and Supplementary Material: Fig. S1). Our results are congruent with a recent meta-analysis where the authors concluded that species inventories of macroinvertebrates obtained with DNA metabarcoding showed pronounced differences to traditional methods, missing some taxa, but at the same time detecting overlooked diversity (Keck et al. 2022).

Despite the differences recorded, the dominant taxonomic groups recovered were common to both methodologies, namely Annelida:Polychaeta, Arthropoda:Crustacea and Mollusca:Bivalvia, which are well known to dominate benthic communities in Portuguese transitional ecosystems (e.g., Sousa et al. 2006, França et al. 2009, Neto et al. 2010, Rodrigues et al. 2011, Lobo et al. 2017b). Exceptionally, Cnidaria:Hydrozoa dominated in the metabarcoding datasets of some sites of the Vouga estuary (10E/06, 09F/04, 09F/08) (Fig. 5). Many of these species have polyp stages and can experience easy fragmentation of their body or tissues, which can be easily detected through metabarcoding, but go unnoticed through the morphological approach, as already above-mentioned.

### 4.2. The use of a multi-marker strategy increased recovered species through metabarcoding

Overall, only 35 species were identified by both methodologies, out of a global sum of 154 recorded in this study. However, this number would be even lower if only one genetic marker was employed, instead of two. For instance, from the 35 species detected with both approaches, 17 were recovered exclusively in the COI dataset and 7 in the 18S dataset (Fig. 3). In a recent review where 90 publications where analysed (Duarte et al. 2021), partial segments of the COI gene have been by far the most used for targeting benthic invertebrate communities in marine ecosystems, including transitional ecosystems, while 18S has been less used, whereas the concurrent use of both markers is less common (e.g. Cowart et al. 2015, Wangensteen et al. 2018a,b, Leite et al. 2021).

The COI marker is by far the marker for which a higher coverage exists in genetic databases, in particular for dominant groups of benthic marine invertebrates (40 to 80%, for the AMBI checklist, Weigand et al. 2019; 16 to 53%, for Mollusca, Crustacea and Polychaeta occurring in Atlantic Iberia, Leite et al. 2020). While species level resolution can be substantially higher using COI, the primer binding sites can be highly variable (see more discussion about this issue in the next sub-section) failing to anneal with DNA templates from species for which the primers have a lower affinity. On the other hand, 18S has been reported to have low variability in primer binding sites (Tang et al. 2012, Brown et al. 2015), but can lack resolution in species level detections (Cowart et al. 2015) and may amplify many small sized species (< 1mm), such as harpacticoid copepods, ostracods and gastrotrichs, that are not included in current morphology-based bioassessments (Lejzerowicz et al. 2015). However, in our study both markers frequently complemented each other, with benthic invertebrate species, which were representative of different ecological groups and present in the morphology dataset, detected exclusively by 18S: the polychaetes *Capitella capitata* (V), *Eumida sanguinea* (II) and *Nephtys incisa* (II) and the bivalves *Cerastoderma edule* (III), *Mytilus edulis* (III), *Pharus legumen* (I) and *Spisula subtruncata* (I). Although reference sequences exist for all of them in BOLD, these species would have gone unnoticed through metabarcoding if only COI was employed, which can have important ecological implications. Our results are consistent with previous findings for marine invertebrate communities, where a multi-marker strategy can significantly improve the number of recovered species (Wangesteen et al. 2018a,b, Leite et al. 2021), therefore contributing to increase the comprehensiveness and reliability of benthic ecosystems assessments using DNA metabarcoding.

### 4.3. Probable reasons for the failed detection of benthic invertebrates through metabarcoding

Given that 55 invertebrate species were detected exclusively through morphology, we investigated the reasons for this in greater detail. We have found straightforward reasons for failed detection through metabarcoding for most of the species: i) 18 species (20%) do not have DNA sequences on BOLD (COI) and/or SILVA (18S); ii) 2 species (∼2%) had less than 9 reads (the minimum threshold set in the bioinformatic pipeline) (*Asterias rubens* and *Mactra stultorum*) and iii) 26 species (29%) were not recorded in the replicate that was also used for metabarcoding (i.e., R3 of each sampling site), but were identified through morphology in the remaining replicates; hence they were counted as species detected through morphology although they did not occur in the replicate used for DNA metabarcoding (Fig. 6).

For 9 species (ca. 10%) we did not find any apparent reason for failed molecular detection, namely the amphipods *Ampelisca brevicornis*, and *Microdeutopus chelifer*, the isopods *Cyathura carinata, Sphaeroma serratum*, and *Eurydice pulchra*, the decapod *Liocarcinus holsatus*, the bivalves *Fabulina fabula* and *Scrobicularia plana* and the polychaete *Diopatra marocensis* (Supplementary Material: Table S5). All of them are represented with COI sequences in BOLD, and 7 of them with 18S sequences in SILVA. In addition, several specimens of each of these species were present in the analysed samples, with the exception of *D. marocensis, E. pulchra, L. holsatus* and *S. plana* which were represented by only one specimen. However, the robustness of metabarcoding has been well demonstrated in several studies, where sequences from species represented by only one specimen (Lobo et al. 2017b) or from single larvae spiked in environmental samples (Pochon et al. 2013, Zhan et al. 2013), have been detected despite the co-occurrence of a large array of other species. Thus, we consider that the most possible reasons for the absence of these species from the metabarcoding dataset may either include inefficient DNA extraction or mismatches between the selected primers and target templates. Another possibility is occurrence of cryptic diversity, which is common in marine invertebrates (e.g., polychaetes: Lobo et al. 2016, Teixeira et al. 2020; amphipods: Lobo et al. 2017a, Vieira et al. 2020; gastropods: Borges et al. 2016), meaning that the species is thought to be present in the reference libraries, but the specimens collected belong to a divergent lineage that is not represented in the library. They would be identified through morphology, but would be missed by metabarcoding. However, we investigated this possibility and found that reference sequences were all from specimens collected in the region, therefore the chances of missing a cryptic lineage are low.

Although, morphological traits have been reported to influence DNA extraction of marine invertebrates (e.g., body size, presence of chitine or CaCO_3_), in particular from preservative ethanol, in bulk DNA samples species detection does not seem to be much affected (Derycke et al. 2021). In addition, we used a non-destructive method for DNA extraction, involving the temporary immersion of the bulk specimens in an extraction buffer without previous homogenization (Carew et al. 2018, Leite et al. 2021), since homogenization of bulk samples previously to DNA extraction may favour the amplification of non-target taxa (e.g., non-invertebrate metazoans, fungi, protists, bacteria) (Lejzerowicz et al. 2015, Aylagas et al. 2018). From the initial raw reads, 55% and 39% matched with sequences from marine invertebrate species, for COI and 18S, respectively, but, when considering only quality-filtered reads, these percentages increased to as high as 86% and 67%. These rates compare well with other studies, so we may conclude that there was a relatively high success in recovering sequences of the target group (invertebrates) (e.g., up to 66.5% of COI quality-filtered reads, in the study of Aylagas et al. 2018).

Although we cannot completely discard an inefficient DNA extraction, the most possible explanation is indeed possible mismatches that may exist between the primers used and target templates. Marine invertebrate communities dominating benthic estuarine ecosystems are very complex and highly diverse, belonging to phylogenetically distant taxonomic groups (e.g., Annelida, Mollusca, Crustacea) (Lobo et al. 2013, Zhang et al. 2021). Thus, primers used in the PCR reaction may have higher affinity for DNA templates of particular species, which will be preferentially detected compared to other species also present in the sample. In a recent study, DNA metabarcoding was also unable to detect 19 out of 57 morphospecies, for the best performing primer set, and the authors concluded that the most probable reason was indeed the absence of match between the primers employed and the species present in the samples (Derycke et al. 2021). Interestingly, in previous studies, where species were identified in morphology-based assessments and the same primers were used for COI (Lobo et al. 2017b, Leite 2021), no reads were also generated for *Cyathura carinata* in the metabarcoding datasets. In addition, Lobo and co-authors (2017) used four different primers for the COI region, including one pair targeting the complete barcode region and with which sequences from these species have been successfully generated previously (Lobo et al. 2013). In another study using mock communities, *Cyathura carinata* was detected by a single read for a single primer set out of 5 tested sets (Hollatz et al. 2017). However, for the remaining 8 species we did not find the same evidence in previous studies using the same primers (Hollatz et al. 2017, Lobo et al. 2017b, Derycke et al. 2021, Leite 2021). Species which appear to be particularly recalcitrant to metabarcoding should be signalled and their detection success carefully examined in future studies, as it can lead to systematic false negatives in metabarcoding-based biomonitoring. Possible recalcitrant species are one additional reason by which a multi-marker approach may be a better solution to recover as much as possible the diversity in benthic invertebrate samples using DNA metabarcoding. Given that PCR-free approaches are still too expensive (e.g., Dowle et al. 2016, Giebner et al. 2020), the design of primers customized to specific taxonomic groups (e.g., dominant groups such as Mollusca, Annelida, and Arthropoda), may provide an alternative to avoid biases of broad-coverage primers (Westfall et al. 2020).

### 4.4. Morphological and metabarcoding-based indices did not match completely in all sites, but both datasets responded similarly to the environmental gradient

AMBI indices based on presence-absence of species calculated using morphology and metabarcoding-based taxonomic assessments did not match completely in all sites, contrarily to the observed in previous studies (Aylagas et al. 2016, 2018, Lobo et al. 2017b). The most probable reason for this were the differences found, between both methodologies, in the percentage of contribution of each ecological group recovered (Fig. 7). While the majority of the species detected belong to ecological group III, indicating dominance of species tolerant to pollution, an increase in the % of ecological groups I (sensitive species) and II (indifferent species), was observed in metabarcoding-based assessments. The higher resolution of metabarcoding, providing species level identifications for smaller organisms or difficult taxa in morphology-based assessments, such as Platyhelminthes (Group II), Bryozoa (Group II) and Cnidaria (Group I and II), may have contributed to this outcome. Because of this, 6 sites were evaluated with a better ecological condition with metabarcoding-based assessments in Minho and Vouga (02D/05, 02D/04, 02E/07 and 10E/05, 10E/06 and 09F/04), while the opposite was rarely found, with the exception of one site in Lima, which has switched from undisturbed to slightly disturbed (03D/06). Nevertheless, it must be stressed that for most sites (13), the same degree of perturbation (slightly disturbed) was found with both methodologies (Fig. 7). A recent meta-analysis concluded that, the taxonomic inferences for macroinvertebrates derived from the two methods can be very different, with metabarcoding complementing rather than providing identical estimates compared to traditional approaches (Keck et al. 2022). Our results seem to support this conclusion. Whereas traditional morphology-based identification will specifically target macroinvertebrate taxa higher than 0.5 or 1 mm, metabarcoding will recover a higher diversity of organisms, including body and tissue fragments and early developmental stages unnoticed by the traditional approach. Also, smaller taxonomic groups that can be sensitive to environmental stressors, but that are largely ignored in current biomonitoring due to the inherent difficulties in the morphological identification (e.g., prokaryotes, protists, metazoan meiofauna) can be also detected. In addition, for 4 species (*Corophium multisetosum, Crangon crangon, Hediste diversicolor* and *Tritia reticulata*) we were able to detect multiple molecular operational units (MOTUs) in the COI metabarcoding dataset (data not shown, since we worked at species level), and some with maximum genetic divergences as high as 18%. For instance, for *Hediste diversicolor*, 23 different MOTUs were found distributed among the 20 estuarine sites (data not shown). To what extent these different genetic lineages will have differential sensitivities to pollution and environmental stress remains to be tested, and more benchmarking is definitely needed, with as many samples as possible collected under different environmental contexts. This would suggest the interesting possibility that through the use a broader set of informative taxa in the biotic indexes, metabarcoding can potentially augment the resolution of bioassessments enabling a better discrimination among sites, giving a more holistic vision of an entire ecosystem, that otherwise would be considered identical (Pawlowski et al. 2018). Other possibilities would be to develop new indices based on new indicator groups, as developed, for example, for bacteria (e.g., microgAMBI, Aylagas et al. 2017, Borja 2018) or to use taxonomy-free approaches and machine learning predictive models (Apothéloz-Perret-Gentil et al. 2017, Cordier et al. 2017).

Interestingly, despite the differences found in our study, the outcome of CCA analyses was somehow similar for morphology and metabarcoding-based assessments, and also when combining both datasets (Fig. 8). Salinity (and consequently conductivity and pH) was the variable that better explained the differences in benthic invertebrates’ composition among sites, which was patent in all diversity assessments (Supplementary Material: Table S7). In previous assessments conducted in these estuaries, salinity has been pointed out to be among the factors that most clearly influence benthic assemblages (Minho: Sousa et al. 2008; Lima: Sousa et al. 2007; Vouga: Rodrigues et al. 2011; Mondego: Teixeira et al. 2008a,b). Nitrates, which changed in the opposite direction to the salinity gradient, displaying higher values at oligohaline sites (2 to 6 times higher in Lima and 13 to 24 times higher in Mondego), also influenced community structure assessed with both methods (more in the morphology dataset), but this was not reflected in the AMBI indices. It has been reported that in estuarine ecosystems the natural variability, as well as natural events (e.g., extreme climatic events), play an important role in the response of biotic indices (Chainho et al. 2007, Teixeira et al. 2008b, Neto et al. 2010). In a previous study conducted in Mondego estuary, both natural and anthropogenic variability were satisfactorily detected, but only when accounting the information provided by three different indices (AMBI, Margalef and Shannon-Wiener) (Teixeira et al. 2008b). Still, a great proportion of the variability remained unexplained in the CCA analyses (>50%, for all datasets). Sediment features, are recognized as another highly important factor influencing benthic assemblages’ composition and species abundance within salinity zones (Sousa et al. 2007, Teixeira et al. 2008b). Part of this unexplained variation may therefore derive from the specific sediment characteristics within each site, but which were not assessed in this specific APA’s biomonitoring campaign.

## 5. Conclusions

In conclusion, our results support that metabarcoding provide higher estimates of diversity than the morphology-based approach, and the use of a multi-locus strategy increased recovered diversity through metabarcoding. In addition, sequence gaps on genetic databases, but also PCR failure seem to be the main reasons for the absence of species detection in the metabarcoding dataset. Although morphological and metabarcoding-based indices did not match completely in all sites, similar responses to the environmental gradient were obtained with both methods. Thus, our results support that rather than moving towards a DNA-based approach independent of morphology-based methods, a harmonized approach should be employed, where, when possible, both methods should be integrated to complement each other, in order to improve and expedite benthic monitoring. Specimen identification based on morphological taxonomy continues to be invaluable, providing the prime foundation in all biodiversity assessments, importantly enabling an estimation of organisms’ abundances and assessment of prevalent life stages, which is actually conducted in 1 to 2 events per 6-year management cycle (Hering et al. 2010). On the other hand, DNA-based monitoring can be less expensive and more responsive to immediate regulatory and management needs, such as the required for monitoring pollution events or restoration activities, which can be employed in a yearly basis or even two-times/year and possibly with higher spatial density. We anticipate that metabarcoding can also increase the quality of the assessments (representativeness and precision), allowing identifications of all specimens in a sample including larval stages and juveniles, but also small organisms from taxonomic groups that cannot be identified to species level using the traditional approach and that are largely ignored in routine biomonitoring and may be sensitive as well to environmental stress (e.g., nematodes, ciliates, foraminifera), as the current targeted BQEs. In addition, cryptic lineages can also be detected, as we were able to do so for four common bioindicator species, promising a greater taxonomic resolution and improvement of the delineation of tolerance/sensitivity groups commonly used in biotic indices, such as AMBI. DNA-based monitoring can also generate standardized data more amenable to audit and less vulnerable to variability in taxonomic expertise among studies, facilitating direct comparison among independent diversity assessments and that can be more easily articulated at regional, European and even at global scale.

## Supporting information

All supplementary material (Tables + figures)

## Acknowledgments

This work was funded by the project “River2Ocean – Socio-ecological and biotechnological solutions for the conservation and valorization of aquatic biodiversity in the Minho Region” (NORTE-01-0145-FEDER-000068), co-financed by the European Regional Development Fund (ERDF), through Programa Operacional Regional do Norte (NORTE 2020), by the “Contrato-Programa” UIDB/04050/2020 funded by national funds through the FCT I.P (Foundation for Science and Technology) and the project MESCLA – “Melhorar e Complementar os Critérios de Classificação do Estado das Massas de Água de Transição e Costeiras” (POSEUR-03-2013-FC-000001). Financial support granted by the FCT to SD (CEECIND/00667/2017), BRL (PD/BD/127994/2016), MALT (SFRH/BD/131527/2017), and PEV (through the project NIS-DNA: PTDC/BIA-BMA/29754/2017) is also acknowledged. The authors also want to thank to Àngel Borja for helping in the decision of the ecological categories of species that are not yet contemplated in the AMBI list.

## References

Apothéloz-Perret-Gentil L, Cordonier A, Straub F, Iseli J, Esling P, Pawlowski J (2017) Taxonomy-free molecular diatom index for high-throughput eDNA biomonitoring. Molecular Ecology Resources 17: 1231–1242. https://doi.org/10.1111/1755-0998.12668

Aylagas E, Borja A, Rodríguez-Ezpeleta N (2014) Environmental status assessment using DNA metabarcoding: towards a genetics based marine biotic index (gAMBI). PLoS ONE 9: e90529. https://doi.org/10.1371/journal.pone.0090529

Aylagas E, Borja A, Irigoien X, Rodríguez-Ezpeleta N (2016) Benchmarking DNA metabarcoding for biodiversity-based monitoring and assessment. Frontiers in Marine Science 3: 96. https://doi.org/10.3389/fmars.2016.00096

Aylagas E, Borja A, Muxika I, Rodríguez-Ezpeleta N (2018) Adapting metabarcoding-based benthic biomonitoring into routine marine ecological status assessment networks. Ecological Indicators 95: 194–202. https://doi.org/10.1016/j.ecolind.2018.07.044

Aylagas E, Borja Á, Tangherlini M, Dell’Anno A, Corinaldesi C, Michell CT, Irigoien X, Danovaro R., Rodríguez-Ezpeleta N (2017) A bacterial community-based index to assess the ecological status of estuarine and coastal environments. Marine Pollution Bulletin 114: 679–688. https://doi.org/10.1016/j.marpolbul.2016.10.050

Bacouillard L, Baux N, Dauvin JC, Desroy N, Geiger KJ, Gentil F, Thiébaut É (2020) Long-term spatio-temporal changes of the muddy fine sand benthic community of the Bay of Seine (eastern English Channel). Marine Environmental Research 161: 105062. https://doi.org/10.1016/j.marenvres.2020.105062

Borges LMS, Hollatz C, Lobo J, Cunha AM, Vilela AP, Calado G, Coelho R, Costa AC, Ferreira MSG, Costa MH, Costa FO (2016) With a little help from DNA barcoding: investigating the diversity of Gastropoda from the Portuguese coast. Scientific Reports 6: 20226. https://doi.org/10.1038/srep20226

Borja A (2018) Testing the efficiency of a bacterial community-based index (microgAMBI) to assess distinct impact sources in six locations around the world. Ecological Indicators 85: 594–602. https://doi.org/10.1016/j.ecolind.2017.11.018

Borja A, Franco J, Pérez V (2000). A marine biotic index to establish the ecological quality of soft-bottom benthos within European estuarine and coastal environments. Marine Pollution Bulletin 40: 1100–1114. https://doi.org/10.1016/S0025-326X(00)00061-8

Borja A, Barbone E, Basset A, Borgersen G, Brkljacic M, Elliott M, Garmendia JM, Marques JC, Mazik K, Muxika I, Neto JM, Norling K, Rodríguez JG, Rosati I, Rygg B, Teixeira H, Trayanova A (2011) Response of single benthic metrics and multi-metric methods to anthropogenic pressure gradients, in five distinct European coastal and transitional ecosystems. Marine Pollution Bulletin 63: 499–513. https://doi.org/10.1016/j.marpolbul.2010.12.009

Brown EA, Chain FJJ, Crease TJ, MacIsaac HJ, Cristescu ME (2015) Divergence thresholds and divergent biodiversity estimates: can metabarcoding reliably describe zooplankton communities? Ecology and Evolution 5: 2234–2251. https://doi.org/10.1002/ece3.1485

Cahill AE, Pearman JK, Borja A, Carugati L, Carvalho S, Danovaro R, Dashfield S, Romain D, Féral J-P, Olenin S, Šiaulys A, Somerfield PJ, Trayanova A, Uyarra MC, Chenuil A (2018) A comparative analysis of metabarcoding and morphology-based identification of benthic communities across different regional seas. Ecology and Evolution 8: 8908–8920. https://doi.org/10.1002/ece3.4283

Camacho C, Coulouris G, Avagyan V, Ma N, Papadopoulos J, Bealer K, Madden TL (2009) BLAST+: architecture and applications. BMC Bioinformatics 10: 421. https://doi.org/10.1186/1471-2105-10-421

Campbell A, Nicholls J (2008) Fauna e Flora do litoral de Portugal e Europa. FAPAS - Fundo Para a Protecção dos Animais Selvagens, 320 pp.

Carew ME, Kellar CR, Pettigrove VJ, Hoffmann AA (2018) Can high-throughput sequencing detect macroinvertebrate diversity for routine monitoring of an urban river? Ecological Indicators 85: 440–450. https://doi.org/10.1016/j.ecolind.2017.11.002

Chainho P, Costa JL, Chaves ML, Dauer DM, Costa MJ (2007) Influence of seasonal variability in benthic invertebrate community structure on the use of biotic indices to assess the ecological status of a Portuguese estuary. Marine Pollution Bulletin 54: 1586–1597. https://doi.org/10.1016/j.marpolbul.2007.06.009

Chariton AA, Stephenson S, Morgan MJ, Steven ADL, Colloff MJ, Court LN, Hardy CM (2015) Metabarcoding of benthic eukaryote communities predicts the ecological condition of estuaries. Environmental Pollution 203: 165–174. http://dx.doi.org/10.1016/j.envpol.2015.03.047

Cordier T, Esling P, Lejzerowicz F, Visco J, Ouadahi A, Martins C, Cedhagen T, Pawlowski J (2017) Predicting the ecological quality status of marine environments from eDNA metabarcoding data using supervised machine learning. Environmental Science and Technology 51: 9118–9126. https://doi.org/10.1021/acs.est.7b01518

Comeau AM, Douglas GM, Langille MGI (2017) Microbiome helper: a custom and streamlined workflow for Microbiome Research. mSystems 2: e00127–16 https://doi.org/10.1128/msystems.00127-16

Cowart DA, Pinheiro M, Mouchel O, Maguer M, Grall J, Miné J, Arnaud-Haond S (2015) Metabarcoding is powerful yet still blind: a comparative analysis of morphological and molecular surveys of seagrass communities. PLoS ONE 10: e0117562. https://doi.org/10.1371/journal.pone.0117562

Cristescu ME (2014) From barcoding single individuals to metabarcoding biological communities: towards an integrative approach to the study of global biodiversity. Trends in Ecology and Evolution 29: 566–571. https://doi.org/10.1016/j.tree.2014.08.001

Derycke S, Maes S, Van den Bulcke L, Vanhollebeke J, Wittoeck J, Hillewaert H, Ampe B, Haegeman A, Hostens K, De Backer A (2021) Detection of macrobenthos species with metabarcoding is consistent in bulk DNA but dependent on body size and sclerotization in eDNA from the ethanol preservative. Frontiers in Marine Science 8: 637858. https://doi.org/10.3389/fmars.2021.637858

Dowle EJ, Pochon X, Banks JC, Shearer K, Wood SA (2016) Targeted gene enrichment and high-throughput sequencing for environmental biomonitoring: a case study using freshwater macroinvertebrates. Molecular Ecology Resources 16: 1240–1254. https://doi.org/10.1111/1755-0998.12488

Duarte S, Leite BR, Feio MJ, Costa FO, Filipe AF (2021) Integration of DNA-based approaches in aquatic ecological assessment using benthic macroinvertebrates. Water 13: 331. https://doi.org/10.3390/w13030331

European Commission (2000) Directive 2000/60/EC of the European Parliament and of the Council of 23 October 2000 establishing a framework for Community action in the field of water policy. https://eur-lex.europa.eu/eli/dir/2000/60/oj

European Commission (2008) Directive 2008/56/EC of the European Parliament and of the Council of 17 June 2008 establishing a framework for community action in the field of marine environmental policy (Marine Strategy Framework Directive). https://eur-lex.europa.eu/eli/dir/2008/56/oj

European Commission (2011) Common Implementation Strategy for the Water Framework Directive (2000/60/EC). Guidance document No. 14 – Guidance document on the Intercalibration Process 2008-2011. https://circabc.europa.eu/ui/group/9ab5926d-bed4-4322-9aa7-9964bbe8312d/library/1bfd8ec3-bac3-4ea0-9d8c-e70c0f9ea3db/details

Fais M, Duarte S, Vieira PE, Sousa R, Hajibabaei M, Canchaya CA, Costa FO (2020) Small-scale spatial variation of meiofaunal communities in Lima estuary (NW Portugal) assessed through metabarcoding. Estuarine, Coastal and Shelf Science. 238: 106683. https://doi.org/10.1016/j.ecss.2020.106683

França S, Vinagre C, Pardal MA, Cabral HN (2009) Spatial and temporal patterns of benthic invertebrates in the Tagus estuary, Portugal: comparison between subtidal and an intertidal mudflat. Scientia Marina 73: 307–318. https://doi.org/10.3989/scimar2009.73n2307

Fontes JT, Vieira PE, Ekrem T, Soares P, Costa FO (2021) BAGS: an automated Barcode, Audit & Grade System for DNA barcode reference libraries. Molecular Ecology Resources 21: 573–583. https://doi.org/10.1111/1755-0998.13262

Giebner H, Langen K, Bourlat SJ, Kukowka S, Mayer C, Astrin JJ, Misof B, Fonseca VG (2020) Comparing diversity levels in environmental samples: DNA sequence capture and metabarcoding approaches using 18S and COI genes. Molecular Ecology Resources 20: 1333–1345. https://doi.org/10.1111/1755-0998.13201

Hajibabaei M (2012) The golden age of DNA metasystematics. Trends in Genetics 28: 535–537. https://doi.org/10.1016/j.tig.2012.08.001

Hammer Ø, Harper DAT, Ryan PD (2001) PAST: Paleontological statistics software package for education and data analysis. Palaeontologia Electronica 4, 9 pp. http://palaeo-electronica.org/2001_1/past/issue1_01.htm

Hayward PJ, Nelson-Smith T, Shields C (1996). Collins pocket guide – sea shore of Britain & Europe. HarperCollins Publishers, 352 pp.

Hayward PJ, Ryland JS (2017) Handbook of the marine fauna of North-West Europe. Oxford University Press. https://doi.org/10.1093/acprof:oso/9780199549443.001.0001

Hering D, Borja A, Carstensen J, Carvalho L, Elliott M, Feld CK, Heiskanen A-S, Johnson RK, Moe J, Pont D, Solheim AL, van de Bund W (2010) The European Water Framework Directive at the age of 10: a critical review of the achievements with recommendations for the future. Science of the Total Environment 408: 4007–4019. https://doi.org/10.1016/j.scitotenv.2010.05.031

Hering D, Borja A, Jones JI, Pont D, Boets P, Bouchez A, Bruce K, Drakare S, Hänfling B, Kahlert M, Leese F, Meissner K, Mergen P, Reyjol Y, Segurado P, Vogler A, Kelly M (2018) Implementation options for DNA-based identification into ecological status assessment under the European Water Framework Directive. Water Research 138: 192–205. https://doi.org/10.1016/j.watres.2018.03.003

Hollatz C, Leite BR, Lobo J, Froufe H, Egas C, Costa FO (2017) Priming of a DNA metabarcoding approach for species identification and inventory in marine macrobenthic communities. Genome 60: 260–271. https://doi.org/10.1139/gen-2015-0220

Houba VJG, Novozamsky I, Uittenbogaard J, Lee JJ van der (1987) Automatic determination of “total soluble nitrogen” in soil extracts. Landwirtschaftliche Forschung 40: 295–302.

Illumina (2013) 16S Metagenomic sequencing library preparation manual - preparing 16S ribosomal RNA gene amplicons for the Illumina MiSeq System, 28 pp.

Ivanova NV, De Waard JR, Hebert PDN (2006) An inexpensive, automation-friendly protocol for recovering high-quality DNA. Molecular Ecology Notes 6: 998–1002. https://doi.org/10.1111/j.1471-8286.2006.01428.x

Keck F, Blackman RC, Bossart R, Brantschen J, Couton M, Hürlemann S, Kirschner D, Locher N, Zhang H, Altermatt F (2022) Meta-analysis shows both congruence and complementarity of DNA and eDNA metabarcoding to traditional methods for biological community assessment. Molecular Ecology 31: 1820–1835. https://doi.org/10.1111/mec.16364

Kozich JJ, Westcott SL, Baxter NT, Highlander SK, Schloss PD (2013) Development of a dual-index sequencing strategy and curation pipeline for analyzing amplicon sequence data on the MiSeq Illumina sequencing platform. Applied and Environmental Microbiology 79: 5112–5120. https://doi.org/10.1128/AEM.01043-13

Krom MD (1980) Spectrophotometric determination of ammonia: a study of a modified Berthelot reaction using salicylate and dichloroisocyanurate. Analyst 105: 305–316, https://doi.org/10.1039/an9800500305

Kroon H (1993) Determination of nitrogen in water: comparison of a continuous-flow method with on-line UV digestion with the original Kjeldahl method. Analytica Chimica Acta 276: 287–293. https://doi.org/10.1016/0003-2670(93)80396-3

Leese F, Bouchez A, Abarenkov K, Altermatt F, Borja A, Bruce K, Ekrem T, Čiampor Jr F, Ciamporová-Zatovicová Z, Costa FO, Duarte S, Elbrecht V, Fontaneto D, Franc A, Geiger MF, Hering D, Kahlert M, Stroil BK, Kelly M, Keskin E, Liska I, Mergen P, Meissner K, Pawlowski J, Penev L, Reyjol Y, Rotter A, Steinke D, van der Wal B, Vitecek S, Zimmermann J, Weigand AM (2018) Chapter Two - Why we need sustainable networks bridging countries, disciplines, cultures and generations for aquatic biomonitoring 2.0: a perspective derived from the DNAqua-Net COST Action. Advances in Ecological Research 58: 63–99. https://doi.org/10.1016/bs.aecr.2018.01.001

Leite BR (2021) Monitoring coastal benthic colonization of artificial substrates with the support of DNA metabarcoding approaches. PhD thesis in Biology Specialization in Integrated Management of the Sea, University of Minho, Braga, Portugal, 212 pp.

Leite BR, Vieira PE, Teixeira MAL, Lobo-Arteaga J, Hollatz C, Borges LMS, Duarte S, Troncoso JS, Costa FO (2020) Gap-analysis and annotated reference library for supporting macroinvertebrate metabarcoding in Atlantic Iberia. Regional Studies in Marine Science 36: 101307. https://doi.org/10.1016/j.rsma.2020.101307

Leite BR, Vieira PE, Troncoso JS, Costa FO (2021) Comparing species detection success between molecular markers in DNA metabarcoding of coastal macroinvertebrates. Metabarcoding and Metagenomics 5: e70063. https://doi.org/10.3897/mbmg.5.70063

Lejzerowicz F, Esling P, Pillet L, Wilding TA, Black KD, Pawlowski J (2015) High-throughput sequencing and morphology perform equally well for benthic monitoring of marine ecosystems. Scientific Reports 5: 13932. https://doi.org/10.1038/srep13932

Leray M, Yang JY, Meyer CP, Mills SC, Agudelo N, Ranwez V, Boehm JT, Machida RJ (2013) A new versatile primer set targeting a short fragment of the mitochondrial COI region for metabarcoding metazoan diversity: application for characterizing coral reef fish gut contents. Frontiers in Zoology 10: 34. https://doi.org/10.1186/1742-9994-10-34

Lincoln RJ (1979) British Marine Amphipoda: Gammaridea. British Museum (Natural History).

Lobo J, Costa PM, Teixeira MAL, Ferreira MSG, Costa MH, Costa FO (2013) Enhanced primers for amplification of DNA barcodes from a broad range of marine metazoans. BMC Ecology 13: 34. https://doi.org/10.1186/1472-6785-13-34

Lobo J, Teixeira MAL, Borges LMS, Ferreira MSG, Hollatz C, Gomes PT, Sousa R, Ravara A, Costa MH, Costa FO (2016) Starting a DNA barcode reference library for shallow water polychaetes from the southern European Atlantic coast. Molecular Ecology Resources 16: 298–313. https://doi.org/10.1111/1755-0998.12441

Lobo J, Ferreira MS, Antunes IC, Teixeira MAL, Borges LMS, Sousa R, Gomes PA, Costa MH, Cunha MR, Costa FO (2017a) Contrasting morphological and DNA barcode-suggested species boundaries among shallow-water amphipod fauna from the southern European Atlantic coast. Genome 60: 147–157. https://doi.org/10.1139/gen-2016-0009

Lobo J, Shokralla S, Costa MH, Hajibabaei M, Costa FO (2017b) DNA metabarcoding for high-throughput monitoring of estuarine macrobenthic communities. Scientific Reports 7: 15618. https://doi.org/10.1038/s41598-017-15823-6

Martins FMS, Porto M, Feio MJ, Egeter B, Bonin A, Serra SRQ, Tarbelet P, Beja P (2020) Modelling technical and biological biases in macroinvertebrate community assessment from bulk preservative using multiple metabarcoding markers. Molecular Ecology 30: 3221–3238. https://doi.org/10.1111/mec.15620

Neto JM, Teixeira H, Patrício J, Baeta A, Veríssimo H, Pinto R, Marques JC (2010) The response of estuarine macrobenthic communities to natural-and human-induced changes: dynamics and ecological quality. Estuaries and Coasts 33: 1327–1339. https://doi.org/10.1007/s12237-010-9326-x

Pawlowski J, Kelly-Quinn M, Altermatt F, Apothéloz-Perret-Gentil L, Beja P, Boggero A, Borja A, Bouchez A, Cordier T, Domaizon I, Feio MJ, Filipe AF, Fornaroli R, Graf W, Herder J, van der Hoorn B, Jones JI, Sagova-Mareckova M, Moritz C, Barquín J, Piggott JJ, Pinna M, Rimet F, Rinkevich B, Sousa-Santos C, Specchia V, Trobajo R, Vasselon V, Vitecek S, Zimmerman J, Weigand A, Leese F, Kahlert M (2018) The future of biotic indices in the ecogenomic era: Integrating (e)DNA metabarcoding in biological assessment of aquatic ecosystems. Science of The Total Environment 637-638: 1295–1310. https://doi.org/10.1016/j.scitotenv.2018.05.002

Pochon X, Bott NJ, Smith KF, Wood SA (2013) Evaluating detection limits of Next-Generation Sequencing for the surveillance and monitoring of international marine pests. PLoS ONE 8: e73935. https://doi.org/10.1371/journal.pone.0073935

Pruesse E, Peplies J, Glöckner FO (2012) SINA: accurate high throughput multiple sequence alignment of ribosomal RNA genes. Bioinformatics 28: 1823–1829 https://doi.org/10.1093/bioinformatics/bts252

Quast C, Pruesse E, Yilmaz P, Gerken J, Schweer T, Yarza P, Peplies J, Glöckner FO (2013) The SILVA ribosomal RNA gene database project: improved data processing and web-based tools. Nucleic Acids Research 41: D590–D596. https://doi.org/10.1093/nar/gks1219

Ratnasingham S (2019) mBRAVE: The Multiplex Barcode Research and Visualization Environment. Biodiversity Information Science and Standards 3: e37986. https://doi.org/10.3897/biss.3.37986

Ratnasingham S, Hebert PDN (2007) BOLD: The Barcode of Life Data System. Molecular Ecology Notes 7: 355–364. https://doi.org/10.1111/j.1471-8286.2007.01678.x

Rodrigues AM, Quintino V, Sampaio L, Freitas R, Neves R (2011) Benthic biodiversity patterns in Ria de Aveiro, Western Portugal: environmental-biological relationships. Estuarine, Coastal and Shelf Science 95: 338–348. https://doi.org/10.1016/j.ecss.2011.05.019

Rognes T, Flouri T, Nichols B, Quince C, Mahé F (2016) VSEARCH: a versatile open-source tool for metagenomics. PeerJ 4: e2584. https://doi.org/10.7717/peerj.2584

Rosenberg R, Blomqvist M, Nilsson HC, Cederwall H, Dimming A (2004) Marine quality assessment by use of benthic species-abundance distributions: a proposed new protocol within the European Union Water Framework Directive. Marine Pollution Bulletin 49: 728–739. https://doi.org/10.1016/j.marpolbul.2004.05.013

Salas F, Neto JM, Borja A, Marques JC (2004) Evaluation of the applicability of a marine biotic index to characterize the status of estuarine ecosystems: the case of Mondego estuary (Portugal). Ecological Indicators 4: 215–225. https://doi.org/10.1016/j.ecolind.2004.04.003

Schmieder R, Edwards R (2011) Quality control and preprocessing of metagenomic datasets. Bioinformatics 27: 863–864. https://doi.org/10.1093/bioinformatics/btr026

Schloss PD, Westcott SL, Ryabin T, Hall JR, Hartmann M, Hollister EB, Lesniewski RA, Oakley BB, Parks DH, Robinson CJ, Sahl JW, Stres B, Thallinger GG, Van Horn DJ, Weber CF (2009) Introducing mothur: Open-source, platform-independent, community-supported software for describing and comparing microbial communities. Applied and Environmental Microbiology 75: 7537–7541. https://doi.org/10.1128/AEM.01541-09

Solan M, Cardinale BJ, Downing AL, Engelhardt KAM, Ruesink JL, Srivastava DS (2004) Extinction and ecosystem function in the marine benthos. Science 306: 1177–1180. https://doi.org/10.1126/science.1103960

Sousa R, Dias S, Antunes JC (2006) Spatial subtidal macrobenthic distribution in relation to abiotic conditions in the Lima estuary, NW of Portugal. Hydrobiologia 559: 135–148. https://doi.org/10.1007/s10750-005-1371-2

Sousa R, Dias S, Antunes C (2007) Subtidal macrobenthic structure in the lower Lima estuary, NW of Iberian Peninsula. Annales Zoologici Fennici 44: 303–313. https://www.jstor.org/stable/23736773

Sousa R, Dias S, Freitas V, Antunes C (2008) Subtidal macrozoobenthic assemblages along the River Minho estuarine gradient (north-west Iberian Peninsula). Aquatic Conservation: Marine and Freshwater Ecosystems 18: 1063–1077. https://doi.org/10.1002/aqc.871

Steinke D, deWaard SL, Sones JE, Ivanova NV, Prosser SWJ, Perez K, Braukmann TWA, Milton M, Zakharov EV, deWaard JR, Ratnasingham S, Hebert PDN (2022) Message in a bottle–Metabarcoding enables biodiversity comparisons across ecoregions. GigaScience 11: giac040. https://doi.org/10.1093/gigascience/giac040

Steyaert M, Priestley V, Osborne O, Herraiz A, Arnold R, Savolainen V (2020) Advances in metabarcoding techniques bring us closer to reliable monitoring of the marine benthos. Journal of Applied Ecology 57: 2234–2245. https://doi.org/10.1111/1365-2664.13729

Stoeck T, Bass D, Nebel M, Christen R, Jones MDM, Breiner H-W, Richards TA (2010) Multiple marker parallel tag environmental DNA sequencing reveals a highly complex eukaryotic community in marine anoxic water. Molecular Ecology 19: 21–31. https://doi.org/10.1111/j.1365-294X.2009.04480.x

Tang CQ, Leasi F, Obertegger U, Kieneke A, Barraclough TG, Fontaneto D (2012) The widely used small subunit 18S rDNA molecule greatly underestimates true diversity in biodiversity surveys of the meiofauna. Proceedings of the National Academy of Sciences 109: 16208–16212. https://doi.org/10.1073/pnas.1209160109

Teixeira H, Salas F, Borja A, Neto JM, Marques JC (2008a) A benthic perspective in assessing the ecological status of estuaries: the case of the Mondego estuary (Portugal). Ecological Indicators 8: 404–416. https://doi.org/10.1016/j.ecolind.2007.02.008

Teixeira H, Salas F, Neto JM, Patrício J, Pinto R, Veríssimo H, García-Charton JA, Marcos C, Pérez-Ruzafa A, Marques JC (2008b) Ecological indices tracking distinct impacts along disturbance-recovery gradients in a temperate NE Atlantic Estuary – Guidance on reference values. Estuarine, Coastal and Shelf Science 80: 130–140. https://doi.org/10.1016/j.ecss.2008.07.017

Teixeira H, Neto JM, Patrício J, Veríssimo H, Pinto R, Salas F, Marques JC (2009) Quality assessment of benthic macroinvertebrates under the scope of WFD using BAT, the Benthic Assessment Tool. Marine Pollution Bulletin 58: 1477–1486. https://doi.org/10.1016/j.marpolbul.2009.06.006

Teixeira MAL, Vieira PE, Pleijel F, Sampieri BR, Ravara A, Costa FO, Nygren A (2020) Molecular and morphometric analyses identify new lineages within a large Eumida (Annelida) species complex. Zoologica Scripta 49: 222–235. https://doi.org/10.1111/zsc.12397

Ter Braak CJF, Verdonschot PFM (1995) Canonical correspondence analysis and related multivariate methods in aquatic ecology. Aquatic Sciences 57: 255–289. https://doi.org/10.1007/BF00877430

Thiébaut E, Cabioch L, Dauvin J-C, Retière C, Gentil F (1997) Spatio-temporal persistence of the Abra alba-Pectinaria koreni muddy-fine sand community of the eastern Bay of Seine. Journal of the Marine Biological Association of the United Kingdom 77: 1165–1185. https://doi.org/10.1017/S0025315400038698

Van den Bulcke L, De Backer A, Ampe B, Maes S, Wittoeck J, Waegeman W, Hostens K, Derycke S (2021) Towards harmonization of DNA metabarcoding for monitoring marine macrobenthos: the effect of technical replicates and pooled DNA extractions on species detection. Metabarcoding and Metagenomics 5: e71107. https://doi.org/10.3897/mbmg.5.71107

Vieira PE, Desiderato A, Holdich DM, Soares P, Creer S, Carvalho GR, Costa FO, Queiroga H (2020) Deep segregation in the open ocean: Macaronesia as an evolutionary hotspot for low dispersal marine invertebrates. Molecular Ecology 28: 1784–1800. https://doi.org/10.1111/mec.15052

Vinagre PA, Pais-Costa AJ, Marques JC, Neto JM (2015) Setting reference conditions for mesohaline and oligohaline macroinvertebrate communities sensu WFD: Helping to define achievable scenarios in basin management plans. Ecological Indicators 56: 171–183. https://doi.org/10.1016/j.ecolind.2015.04.008

Wangensteen OS, Cebrian E, Palacín C, Turon X (2018a) Under the canopy: community-wide effects of invasive algae in marine protected areas revealed by metabarcoding. Marine Pollution Bulletin 127: 54–66. https://doi.org/10.1016/j.marpolbul.2017.11.033

Wangensteen OS, Palacín C, Guardiola M, Turon X (2018b) DNA metabarcoding of littoral hard-bottom communities: high diversity and database gaps revealed by two molecular markers. PeerJ 6: e4705. https://doi.org/10.7717/peerj.4705

Weigand H, Beermann AJ, Čiampor F, Costa FO, Csabai Z, Duarte S, Geiger MF, Grabowski M, Rimet F, Rulik B, Strand M, Szucsich N, Weigand AM, Willassen E, Wyler SA, Bouchez A, Borja A, Ciamporová-Zatovicová Z, Ferreira S, Dijkstra K-DB, Eisendle U, Freyhof J, Gadawski P, Graf W, Haegerbaeumer A, van der Hoorn BB, Japoshvili B, Keresztes L, Keskin E, Leese F, Macher JN, Mamos T, Paz G, Pešic V, Pfannkuchen DM, Pfannkuchen MA, Price BW, Rinkevich B, Teixeira MAL, Várbíró G, Ekrem T (2019) DNA barcode reference libraries for the monitoring of aquatic biota in Europe: Gap-analysis and recommendations for future work. Science of the Total Environment 678: 499–524. https://doi.org/10.1016/j.scitotenv.2019.04.247

Westfall KM, Therriault TW, Abbot CL (2020) A new approach to molecular biosurveillance of invasive species using DNA metabarcoding. Global Change Biology 26: 1012–1022. https://doi.org/10.1111/gcb.14886

WoRMS Editorial Board (2021) World Register of Marine Species. Available from http://www.marinespecies.org at VLIZ. Accessed on 2021-11-23. http://doi.org/10.14284/170

Zhan A, Hulák M, Sylvester F, Huang X, Adebayo AA, Abbott CL, Adamowicz SJ, Heath DD, Cristescu ME, MacIsaac HJ (2013) High sensitivity of 454 pyrosequencing for detection of rare species in aquatic communities. Methods Ecology and Evolution 4: 558–565. https://doi.org/10.1111/2041-210X.12037

Zhang Y, Wang J, Lv M, Gao H, Meng LF, A Y, Seim I, Zhang H, Liu S, Zhang L, Liu X, Xu X, Yang H, MI0K+ consortium, Fan G (2021) Diversity, function and evolution of marine invertebrate genomes. bioRxiv, https://doi.org/10.1101/2021.10.31.465852

